# Legacy Data Confounds Genomics Studies

**DOI:** 10.1101/624908

**Authors:** Luke Anderson-Trocmé, Rick Farouni, Mathieu Bourgey, Yoichiro Kamatani, Koichiro Higasa, Jeong-Sun Seo, Changhoon Kim, Fumihiko Matsuda, Simon Gravel

## Abstract

Recent reports have identified differences in the mutational spectra across human populations. While some of these reports have been replicated in other cohorts, most have been reported only in the 1000 Genomes Project (1kGP) data. While investigating an intriguing putative population stratification within the Japanese population, we identified a previously unreported batch effect leading to spurious mutation calls in the 1kGP data and to the apparent population stratification. Because the 1kGP data is used extensively, we find that the batch effects also lead to incorrect imputation by leading imputation servers and a small number of suspicious GWAS associations. Lower-quality data from the early phases of the 1kGP thus continues to contaminate modern studies in hidden ways. It may be time to retire or upgrade such legacy sequencing data.

## Introduction

### Batch Effects in Aging Reference Cohort Data

The last 5 years have seen a drastic increase in the amount and quality of human genome sequence data. Reference cohorts such as the International HapMap Project (***International HapMap Consortium, 2005***), the 1000 Genomes Project (1kGP)(***1000 Genomes Project Consortium, 2010, 2012***; ***Consortium et al., 2015***), and the Simons Genome Diversity project (SGDP)(***Mallick et al., 2016***), for example, have made thousands of genome sequences publicly available for population and medical genetic analyses. Many more genomes are available indirectly through servers providing imputation services (***McCarthy et al., 2016***) or summary statistics for variant frequency estimation (***Lek et al., 2016***).

The first genomes in the 1kGP were sequenced 10 years ago (***van Dijk et al., 2014***). Since then, sequencing platforms have rapidly improved. The second phase of the 1kGP implemented multiple technological and analytical improvements over its earlier phases (***1000 Genomes Project Consortium, 2012***; ***Consortium et al., 2015***), leading to heterogeneous sample preparations and data quality over the course of the project.

Yet, because of the extraordinary value of freely available data, early data from the 1kGP is still widely used to impute untyped variants, to estimate allele frequencies, and to answer a wide range of medical and evolutionary questions. This raises the question of whether and how such legacy data should be included in contemporary analyses alongside more recent cohorts. Mafessoni et al. recently identified batch effects in the 1kGP by looking for individuals with excess LD among distant variants. (***Mafessoni et al., 2018***). Here we point out how these and additional unreported batch effects in the early phases of the 1kGP lead to incorrect genetic conclusions through population genetic analyses and spurious GWAS associations as a result of imputation using the 1kGP as a reference.

### Mutational Signatures

Different mutagenic processes may preferentially affect different DNA motifs. Certain mutagens in tobacco smoke, for example, have been shown to preferentially bind to certain genomic motifs leading to an excess of G to T transversions (***Pfeifer et al., 2002***; ***Pleasance et al., 2010***). Thus, exposure of populations to different mutational processes can be inferred by considering the DNA context of polymorphism in search of *signatures* of different mutational processes (***Alexandrov et al., 2013***; ***Shiraishi et al., 2015***). Such genome-wide mutational signatures have been used as diagnostic tools for cancers (e.g., ***Alexandrov et al.*** (***2013***); ***Shiraishi et al.*** (***2015***)).

In addition to somatic mutational signatures, there has been recent interest in population variation in germline mutational signatures which can be revealed in large sequencing panels. In 2015, Harris reported 50% more TCC *→* TTC mutations in European populations compared to African populations, and this was replicated in a different cohort in 2017 (***Harris, 2015***; ***Harris and Pritchard, 2017***; ***Mathieson and Reich, 2017***). Strong population enrichments of a mutational signature suggests important genetic or environmental differences in the history of each population (***Harris, 2015***; ***Harris and Pritchard, 2017***). Harris and Pritchard further identified distinct mutational spectra across a range of populations, which were further examined in a recent publication by Aikens et al. (***Harris and Pritchard, 2017***; ***Aikens et al., 2019***).

Another signature, *AC → *CC, has been observed at higher frequency in East Asians compared to Africans in the 1kGP and the SGDP (***Aikens et al., 2019***; ***Harris and Pritchard, 2017***). These two studies also found heterogeneous frequency of this signature among 1kGP Japanese individuals. This heterogeneity is intriguing because differences in germline signatures accumulate over many generations. A systematic difference within the Japanese population would suggest sustained environmental or genetic differences across sub-populations within Japan with little to no gene flow. This observation could not be reproduced in SGDP due to the small number of Japanese samples (***Aikens et al., 2019***). We therefore decided to follow up on this observation by using a newly sequenced dataset of Japanese individuals from Nagahama. While we were unable to reproduce the mutational heterogeneity within the Japanese population, we could trace back the source of the discrepancy to a technical artefact in the 1kGP data. In addition to creating biases in mutational signatures, this artefact leads to spurious imputation results which have found their way in recent publications and online resources.

The results section is organized as follows. We first attempt to reproduce the original signal and identify problematic variants in the JPT cohort from the 1kGP. Next, we expand our analysis to the other populations in the 1kGP and identify lists of variants that show evidence for technical bias. Finally, we investigate how these variants have impacted modern genomics analyses.

## Results

### A peculiar mutational signature in Japan

Harris and Pritchard reported an excess of a 3-mer substitution patterns *AC*→*\*CC in a portion of the Japanese individuals in the 1kGP (***Harris and Pritchard, 2017***). Our initial goal was to determine whether this signature could be explained by population structure or technological error. While trying to follow up on this observation in a larger and more recent Japanese cohort from Nagahama, we did not find this particular signature. When comparing the allele frequencies between the Japanese individuals from the 1kGP and this larger dataset, we observed a number of single nucleotide polymorphisms (SNPs) private to one of the two groups (Figure 1). Given the similarity of the two populations, this strongly suggests a technical difference rather than a population structure effect. These mismatches were maintained despite only considering sites that satisfied strict quality masks and Hardy-Weinberg equilibrium in both cohorts.

**Figure 1.**
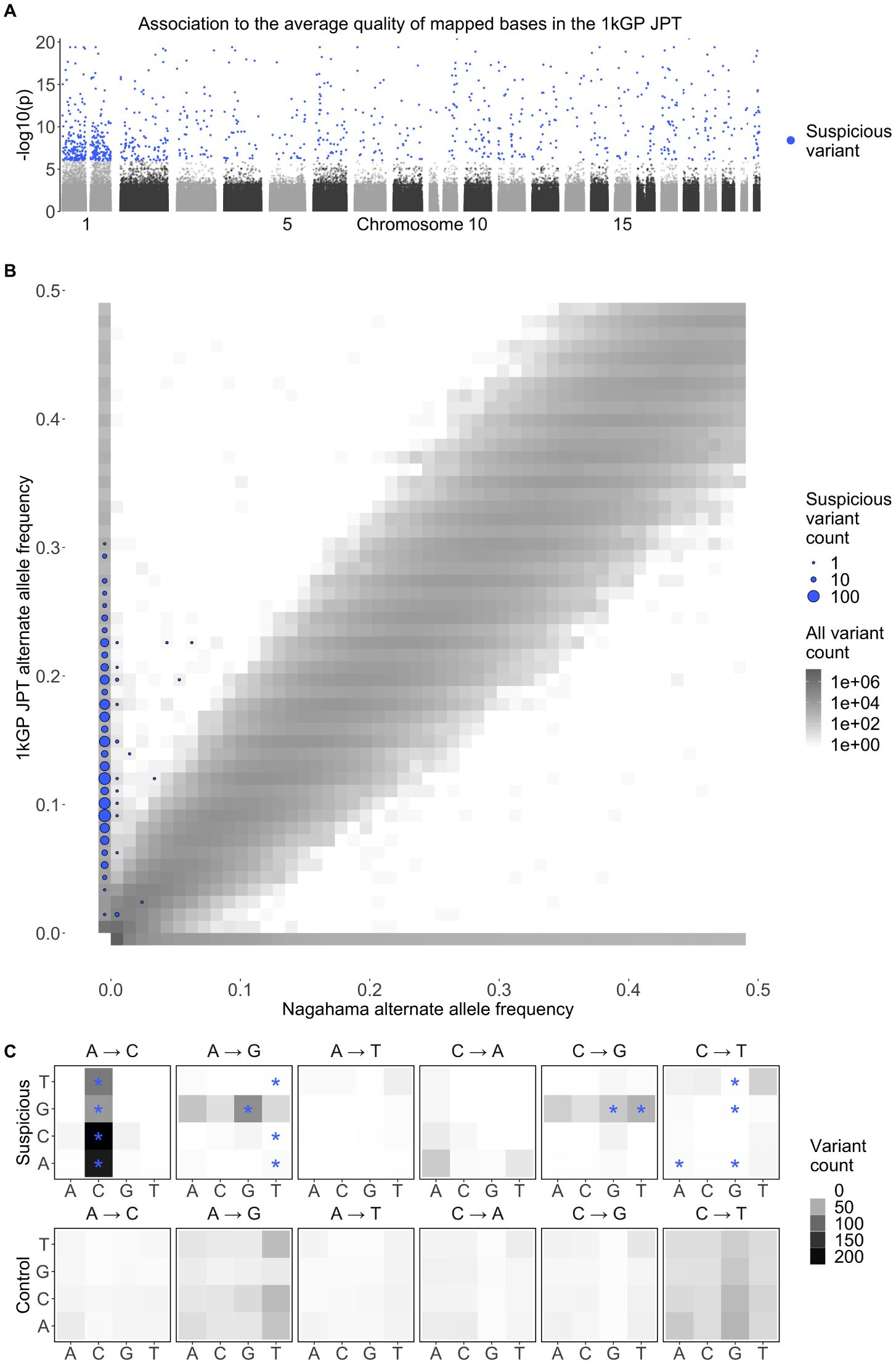
Suspicious mutations carried by individuals with low quality data have distinct mutational profiles, reproduce poorly across studies, and are distributed across the genome. **A** Genome wide association of the average quality of mapped bases *Q* for the 104 Japanese individuals included in the 1000 Genomes Project. This GWAS identified 1034 SNPs associated to the average *Q* of SNPs mapped for an individual, with *p<* 10^−6^ (587 SNPs had *p<* 10^−8^.) **B** Joint frequency spectrum plot of the Japanese from the 1000 Genomes Project and a more recent Japanese dataset from Nagahama. The plot is zoomed in on frequencies below 0.5 for clarity. The size of blue dots are proportional to the number of variants in a given frequency bin that associate with *Q* in the JPT. **C** Mutation spectrum of the 1034 variants that associated with *Q* in the JPT(*p<* 10^−6^), compared to the expectation from the distribution of all SNPs. The majority of the variants with significant associations to *Q* have the *AC*→*\*CC mutational pattern. There is also an enrichment in GA\**→*GG* and GC\**→*GG* mutations. These three enrichments can be summarized as G\*\**→*GG*. Stars (*) indicate a significant deviation from the expected mutational spectrum from all SNPs.

When mismatch sites are removed from the 1kGP data, the *AC*→*\*CC signal disappears (Figure 1). To identify possible technical reasons for the difference, we performed regressions of the prevalence of the *AC*→*\*CC mutational signature against different individual-level quality metrics provided by the 1kGP (see Figure S1). The average quality of mapped bases *Q* per individual stood out as a strong correlate : Individuals with low *Q* show elevated rates of the signature. Thus, sequences called from low-*Q* data contain variants that reproduce poorly across studies and exhibit a particular mutational signature.

To identify SNPs that are likely to reproduce poorly across cohorts without having access to a second cohort, we performed an association study in the JPT for SNPs that associate strongly with low *Q* (Figure 1A). Traditionally, genome wide association studies use genotypes as the independent variable. Here we perform a genotype conditional association test (GCAT), where genotypes are now the dependent variable that we predict using the continuous variable *Q* as the independent variable (***Song et al., 2015***). We use logistic regression of the genotypes on *Q* and identify 587 SNPs with *p<* 10^−8^ and 1034 SNPs with *p<* 10^−6^. While identifying putative low-quality SNPs to exclude, using a higher *p*-value threshold increases the stringency of the filtering (i.e., excluding SNPs with *p <* 10^−6^ is more stringent than excluding SNPS with *p <* 10^−8^). The variants that are associated to *Q* have a significant enrichment in *AC*→*\*CC mutations, GA\**→*GG*, and GC\**→*GG* mutations (Figure 1C). These three enrichments can be summarized as an excess of G\*\**→*GG* in individuals with low *Q*. Statistical significance of these enrichment is computed using a chi-squared test following ***Harris and Pritchard*** (***2017***).

Thus, this mutational signal is heavily enriched in *Q*-associated SNPs, but residual signal remains in non-significant SNPs, presumably because many rare alleles found in individuals with low *Q* remain unidentifiable using association techniques due to lack of power (Figure S2). The removal of individuals with *Q* below 30 successfully removes the *AC*→*\*CC signal enrichment in the Japanese, however other signals identified by Harris and Pritchard appear unchanged including the continental enrichment of *AC*→*\*CC signal in East Asians compared to Africans as reported by Harris and Pritchard and replicated in the SGDP (Figure S3 and S4). For population genetic analyses sensitive to the accumulation of rare variants, the removal of individuals with low *Q* appears preferable to filtering specific low-quality SNPs. For other analyses where quality of imputation matters, identifying *Q*-associated variants may be preferable.

### Identifying suspicious variants in the 1000 Genomes Project

The distribution of *Q* across 1kGP populations shows that many populations have distributions of *Q* scores comparable to that of the JPT, especially populations sequenced in the phase 1 of the project: sequencing done in the early phases of the 1kGP was more variable and overall tended to include lower quality sequencing data (Figure 2 and Figure S5). This variability could result from evolving sequence platform and protocols or variation between sequencing centres. By 2011, older sequencing technologies were phased out, and methods became more consistent, resulting in higher and more uniform quality.

**Figure 2.**
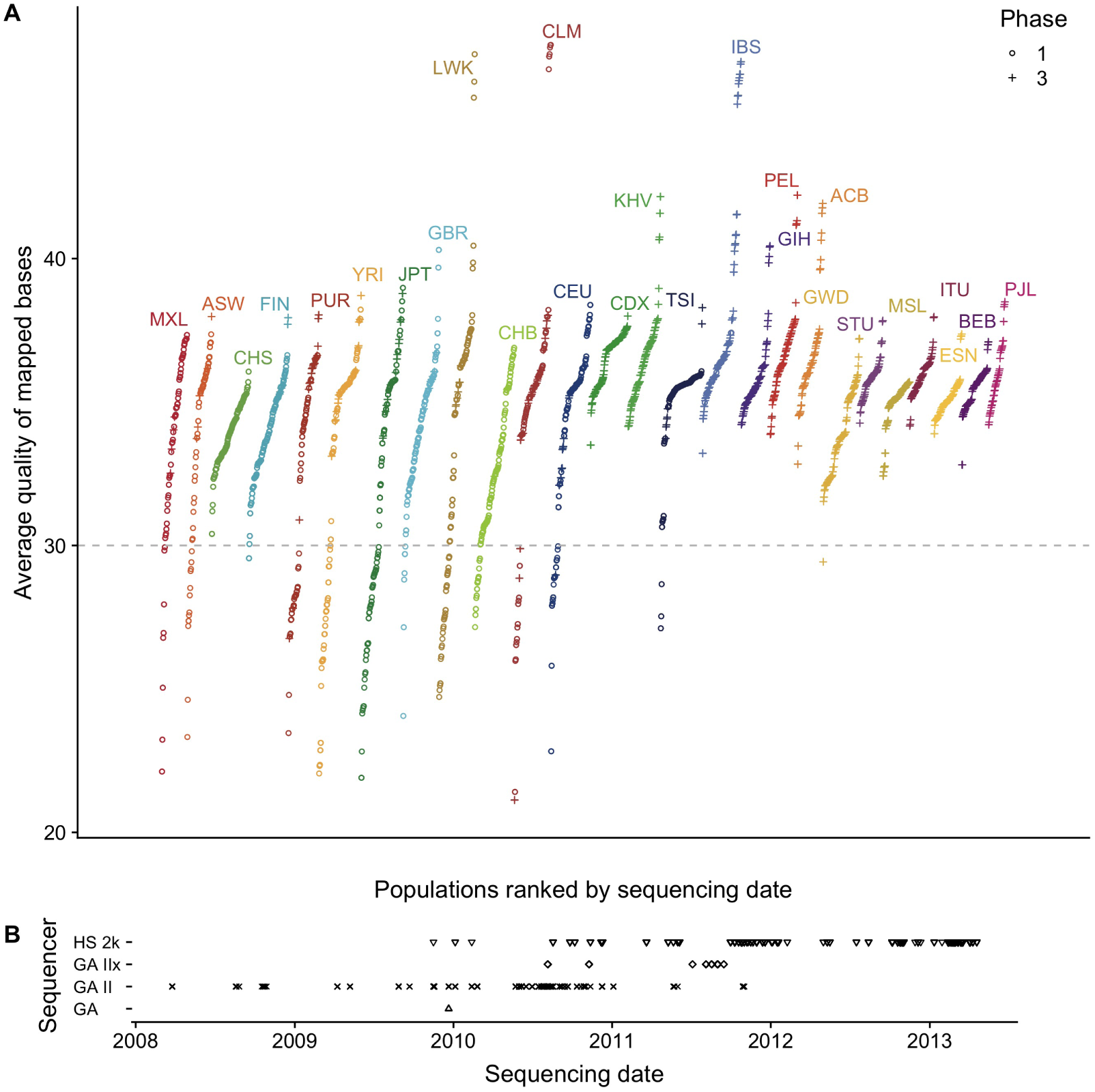
Sampling and sequencing technologies over time in the 1000 Genomes Project. **A** The average quality of mapped bases *Q* for each individual per population included in the 1000 Genomes Project. Populations are ranked by mean sequencing date (the earliest sequencing date was used for individuals with multiple dates). The shape indicates whether the individual was first released in Phase 1 or Phase 3 of the 1000 Genomes project. **B** Sequencing technologies used over the course of the 1000 Genomes Project.

We therefore performed the same reverse GWAS approach in all populations independently, and similarly identified *Q*-associated SNPs in 23 of the 26 populations in the 1kGP, with the phase 1 populations being most affected, with on average four times as many significantly associated sites compared to the phase 3 populations. Over 1,165 variants were independently associated to low *Q* with *p<* 10^−6^ in each (Figure S6).

To build a test statistic to represent the association across all populations simultaneously, we performed a simple logistic regression predicting genotype based on *Q* with the logistic factor analysis (LFA) as an offset to account for population structure or Genotype-Conditional Association Test (GCAT) as proposed by (***Song et al., 2015***). We also considered two alternative approaches to account for confounders, namely using the leading five principal components, and using population membership as covariates. These models were broadly consistent (See Figure S7).

This method identifies a total of 24,390 variants associated to *Q* distributed across the genome with 15,270 passing the 1kGP strict mask filter (Figures S8, S9, S10 and S11). Most analyses below focus on the 15,270 variants satisfying the strict mask, since these variants are unlikely to be filtered by standard pipelines. To account for the large number of tests, we used a two-stage Benjamini & Hochberg step-up FDR-controlling procedure to adjust the p-values using a nominal Type-I error rate *α* = 0.01 (***Benjamini et al., 2006***). We tested SNPs, INDELs and repetitive regions separately as they may have different error rates (Table 1). Lists of *Q*-associated variants and individuals with low *Q* are provided in Supplementary Data.

**Table 1.** Number and proportion of *Q*-associated variants passing the 1000 Genomes Project strict mask per category. Variants that are flagged by the 1000 Genomes Project nested repeat mask file were analyzed separately for FDR calculation. SNPs and INDELs were also analyzed separately. A total of 15,270 are statistically significantly associated to *Q* and pass the 1kGP strict mask. The grey text is the proportion of *Q*-associated variants per category. The number of variants included in the analysis for SNPs, SNPs in repeat regions, INDELs and INDELs in repeat regions are 19,846,786, 6,312,620, 1,770,315 and 586,342 respectively.

|  | Repeat | Proportion | Non-Repeat | Proportion | Total |
| --- | --- | --- | --- | --- | --- |
| SNP | 3,369 | 0.53‰ | 11,059 | 0.56‰ | 14,428 |
| INDEL | 181 | 0.3‰ | 657 | 0.66‰ | 838 |
| Total | 3,550 |  | 11,716 |  | 15,270 |

*Q*-associated variants are distributed across the genome, with chromosome 1 showing an excess of such variants, and other chromosomes being relatively uniform (Figure S12A). Chromosome 1 shows strong enrichments in the *AC*→*\*CC signal compared to other chromosomes despite normalizing for the number of variants tested per chromosome (Figure S12B). At a 1Mb scale, we also see rather uniform distribution with a small number of regions showing an enrichment for such variants (Figure S12C). Three outlying 1Mb regions in chromosomes 1, 2 and 17 have over 30 *Q*-associated variants. Distribution of association statistics in these regions are provided in Figure S13. By contrast, variants that do not pass the 1kGP strict mask are more unevenly distributed across the genome (Figure S12D).

The mutational 3-mer substitution patterns of this list of variants from the GCAT model is similar to the signature identified in the single population test of the 1kGP JPT in that there is an enrichment in *AC*→*\*CC. There is also an enrichment of mutations from TAC, TCT, and TGT to the homonucleotide TTT (Figure S14).

A recent publication by Mafessoni et al. also identified a batch effect in the 1kGP using a method that uses linkage disequilibrium rather than quality metrics to identify 19,196 suspicious variants with 67% of them passing the 1kGP strict mask (***Mafessoni et al., 2018***) (Figure S15A). They identify 17,917 variants significantly associated to abnormal LD patterns that are not associated to *Q*. We find that 1,279 (3%) of the variants they identified are also in our list of suspicious variants and have correlated p-values to those identified using the GCAT method. We also find 23,111 *Q*-associated variants that are not associated to abnormal LD patterns. Interestingly, the variants identified my Mafessoni et al. are not enriched in the mutational spectra described above (Figure S15B). These results indicate that there may a multitude of batch effects in the 1kGP that can only be identified using a suite of association tests.

### Spurious variants, biased genotypes, or cell line artefacts?

To assess whether *Q*-associated variants are spurious variants resulting from sequencing artifacts or real variants exhibiting biased genotyping, we compare the original 1kGP sequence data to more recent sequencing efforts using the same cell lines. *Q*-associated variants that do not reproduce across sequencing experiments are likely the result of sequencing artifacts in the 1kGP. By contrast, *Q*-associated variants that do reproduce across experiments could result from (recurrent) cell line mutations or, more likely, from existing variants whose genotyping depends on *Q*. Finally, given our nominal false discovery rate of *α* = 0.01, we expect approximately 1% of *Q*-associated variants to be false discoveries, i.e., variants that are not associated with *Q* and that should therefore reproduce across experiments.

In 2017, Lan et al. resequenced 83 Han Chinese individuals from the 1kGP (***Lan et al., 2017***). Among the 296 such variants that were *Q*-associated in single-population tests for the CHB or CHS, only 6 are present in the resequenced data (Figure S16). This is slightly more but consistent with the 3 false positives predicted according to the *α* = 0.01 nominal false discovery rate. Thus the majority of *Q*-associated variants in the CHB or CHS appear to be spurious variants.

We did a similar analysis using all variants identified in the GCAT model (rather than only variants significantly associated to *Q* within the CHB and CHS). Of the 15,270 *Q*-associated variants identified globally, 6,307 are polymorphic in the 1kGP for the 83 resequenced individuals (See Figure S17). The vast majority of polymorphisms associated with *Q* are not present at all in the resequencing dataset, suggesting that they are spurious variants. Five variants show differing frequencies between both datasets and are likely explained by biased genotypes. Finally, 1,139 (or 18%) are present in the resequenced data at comparable frequencies. Because this is higher than the false discovery rate, we conclude that the cohort-wide GCAT test identifies predominantly spurious variants, but also variants that have biased genotypes in populations other than the CHB/CHD.

Finally, among the 15,270 *Q*-associated variants, 613 are present on Illumina’s Omni 2.5 chip (See Figure S18). These are likely among the small number of variants that are present in the data but exhibit biased genotyping in 1kGP.

### Suspicious variants impact modern genomics analyses

State of the art imputation servers use a combination of many databases including some that are not freely available. From the perspective of researchers, they act as black-box imputation machines that take observed genotypes as input and return imputed genotypes.

To investigate whether suspicious calls from the 1kGP are imputed into genotyping studies, we submitted genotype data for the first two chromosomes of the 1kGP genotype data to the Michigan Imputation Server. We found that all of the variants associated with *Q* were imputed back in the samples. This suggests that the imputation reference panel still includes individuals with low *Q*, and the dubious variants will be imputed in individuals who most closely match the low-quality individual. These *Q*-associated variants could also compromise the imputation of nearby real-variants, however when considering the imputation scores of genotype data from Japanese individuals from Nagahama, there does not appear to be any impact on nearby variant imputation scores (Figure S19).

We searched the literature for any GWAS that might have reported these *Q*-associated variants as being significantly associated with some biological trait, even though there is no particular reason for these variants to be associated with phenotypes. The NHGRI-EBI Catalog of published genome-wide association studies identified seventeen recent publications that had reported these variants as close to or above the genome-wide significant threshold (Table 2).

**Table 2.** Recent publications that reported *Q*-associated variants as close to or above the genome-wide significant threshold. The variants reaching genome wide significance have a star (*). The black text colour indicates that this variant is twice as frequent in individuals with *Q* < 30, grey text colour indicates that these variants are less than twice as frequent in individuals with *Q* < 30 (See Figure S21). † Attention deficit hyperactivity disorder. ‡ 3-hydroxy-1-methylpropylmercapturic acid.

| Pubmed ID | Disease/Trait | rsID | GWAS<br>-log<sub>10</sub> *p* | *Q*<br>-log<sub>10</sub> *p*<br>(adjusted) |
| --- | --- | --- | --- | --- |
| 28654678 | EBV copy number in lymphoblastoid cell lines | rs201761909 | 5.7 | 78.11 |
|  |  | rs201130852 | 5.05 | 72.28 |
|  |  | rs201255786 | 5.7 | 68.97 |
|  |  | rs200655768 | 6.52 | 66.67 |
|  |  | rs184202621 | 5.52 | 60.45 |
|  |  | rs80274284 | 6 | 56.15 |
|  |  | rs200699422 | 5.3 | 7.43 |
| 23527680 | ADHD<sup>†</sup> | rs6057648 | 5.4 | 20.5 |
| 28928442 | Cold sores | rs201471471 | 6.52 | 7.87 |
| 26053186 | HMPMA<sup>‡</sup> levels in smokers | rs60136336 | 5.7 | 2.25 |
| 28270201 | HDL cholesterol | rs453755 | 7.52 | 5.29 |
| 23023329 | Prostate cancer | rs103294 | *15.3 | 4.32 |
| 28334899 | HDL cholesterol | rs103294 | *29.3 | 4.32 |
| 28240269 | Blood protein levels | rs103294 | *72.7 | 4.32 |
| 27863252 | High light scatter reticulocyte count | rs3794738 | *13.15 | 3.73 |
| 29534301 | Response to hepatitis B vaccine | rs9273062 | *9.7 | 3.36 |
| 21386085 | Metabolic syndrome | rs301 | *10.52 | 3.02 |
| 26830138 | Alzheimer disease and age of onset | rs77894924 | 6.7 | 2.77 |
| 29617998 | Intraocular pressure | rs4963156 | *22.4 | 2.52 |
| 28698626 | Immunoglobulin A vasculitis | rs11015915 | 5.05 | 2.45 |
| 26974007 | Chronic inflammatory diseases | rs3124998 | *8.05 | 2.33 |
| 26634245 | Post bronchodilator FEV1/FVC ratio | rs451000 | 6 | 2.28 |
|  |  | rs443874 | 5.3 | 2.26 |
|  |  | rs400942 | 6 | 2.2 |
| 25918132 | Diisocyanate-induced asthma | rs76780579 | 6 | 2.09 |

Eleven of these studies included the 1kGP in their reference panel for imputation (***Xu et al., 2012***; ***Lutz et al., 2015***; ***Park et al., 2015***; ***Astle et al., 2016***; ***Herold et al., 2016***; ***Suhre et al., 2017***; ***López-Mejías et al., 2017***; ***Tian et al., 2017***; ***Spracklen et al., 2017***; ***Nagy et al., 2017***; ***Gao et al., 2018***) and another used the 1kGP sequence data and cell lines directly (***Mandage et al., 2017***). One study used an in-house reference panel for imputation (***Nishida et al., 2018***), two studies genotyped individuals and imputed the data using the HapMap II as a reference database for imputation (***Kraja et al., 2011***; ***Ebejer et al., 2013***) and two studies used genotyping chip data (***Yucesoy et al., 2015***; ***Ellinghaus et al., 2016***).

All of these articles used a variety of strict quality filters, including Hardy-Weinberg equilibrium test, deviations in expected allele frequency and sequencing data quality thresholds. They also removed rare alleles and alleles with high degrees of missingness. Indeed, we expect a large number of *Q*-associated variants to be filtered out by some quality controls like the Hardy-Weinberg equilibrium test. Even though the studies used state-of-the-art quality controls, the variants were imputed onto genotype data and reached genome wide significance for association with biological traits. However, the fact that some of these variants in other studies are not removed and that the great majority of these variants are missing from higher quality datasets means that these *Q*-associated variants should be flagged for removal to avoid spurious association.

These associations are not necessarily incorrect – a weak but significant bias in imputation may still result in a correct associations. To distinguish between variants with weak but significant association with *Q* from variants with strong biases, we distinguished between variants where the allele frequency difference between individuals with low- and high-*Q* is larger than a factor of two (which naturally separates two clusters of variants on Figure S20). The majority (92.7%) of the *Q*-associated variants are strongly biased in that they are more than twice as frequent in individuals with low-*Q* compared to high-*Q* data. By contrast, most *Q*-associated variants reported in the GWAS catalogue had weak bias (See Figure S21), with three exceptions. One study that reports associations with seven highly biased *Q*-associated variants considered copy number of Epstein-Barr virus (EBV) sequence in the 1kGP as a phenotype (***Mandage et al., 2017***). These seven variants were not flanked by LD peaks and were correctly removed from downstream analyses however, they were still included in the NHGRI-EBI Catalog (***Mandage et al., 2017***). It is plausible that the EBV copy number phenotype is sensitive to mapping and confounded by the same technical artefacts that lead to biased SNP calling.

## Discussion

The variants identified in this study could be explained by technical artifacts from legacy technologies. Different sequencing technologies will have different error profiles. A report comparing the Genome Analyzer II (GAII) to the Illumina HiSeq found that the GAII had much higher rates of reads below a quality score of 30 (***Minoche et al., 2011***) with, for instance, different patterns of quality decrease along reads. Differences in read quality and error profiles in turn require different calling pipelines.

An enrichment in mutational profiles in one chromosome (Figure 12B) is difficult to explain through biological mechanisms or sequencing technology as neither are expected to produce systematic biases across chromosomes. Subtle differences in how these data were integrated or processed could explain such biases, as chromosomes are commonly treated and analyzed separately. However, to pinpoint the precise technical source of the discrepancy would require further forensic inquiries into the details of the heterogeneous sample preparation, combination of sequencing technologies and data processing pipelines used throughout the 1kGP. Given the progress in sequencing and calling that occurred since the early phases of the 1kGP (Figure 2), it is likely that the source of these biases is not longer being actively introduced in recent sequence data.

However, because the 1kGP data is widely used as a reference database, these variants are still being imputed onto new genotype data and can then impact association studies for a variety of phenotypes. Even though significant association of a variant with a quality metric is not in itself an indication that the variant is spurious, we would recommend to carefully examine GWAS associations for such variants, e.g. by repeating the analysis without the 1kGP as part of the imputation panel.

For analyses where individual variants cannot be examined individually (mutation profiles, distributions of allele frequencies, polygenic risk scores), we would recommend to simply discard the individuals with *Q <* 30 and to filter out the LD-associated variants identified by ***Mafessoni et al.*** (***2018***) and the *Q*-associated variants we identified (lists of such variants and sample IDs are provided in the Supplementary Data). We also recommend that imputation servers discard individuals with low *Q* (or at least provide the option of performing the imputation without). Given the value of freely accessible data, resequencing individuals with low *Q* would also likely be a worthwhile investment for the community.

## Conclusion

On a technical front, we were surprised that strong association between variants and technical covariates in the 1kGP project had not been identified before. The genome-wide logistic regression analysis of genotype on quality metric is straightforward, and should probably be a standard in a variety of -omics studies. The logistic factor analysis is more computationally demanding but produces more robust results (***Song et al., 2015***). All three approaches accounting for population structure we tested produce comparable results.

More generally, to improve the quality of genomic reference datasets, we can proceed by addition of new and better data and by better curation of existing data. Given rapid technological progress, the focus of genomic research is naturally on the data generation side. However, cleaning up existing databases is also important to avoid generating spurious results. The present findings suggest that a substantial fraction of data from the final release of the 1kGP project is overdue for retirement or re-sequencing.

## Methods

### Code and data availability

Since this data is primarily performed using publicly available data, we provide fully reproducible and publicly available on GitHub. This repository includes scripts used for data download, processing, analysis and plotting.

### Metadata

The metadata used in this analysis was compiled from each of the index files from the 1kGP file system. Average quality of mapped bases *Q* per sample was obtained from the BAS files associated with each alignment file. Each BAS file has metadata regarding each sequencing event for each sample. If a sample was sequenced more than once, we took the average of each *Q* score from each sequencing instance. The submission dates and sequencing centres for each sample in the analysis was available in the sequence index files.

### Quality Controls

For the mutation spectrum analysis, we reproduced the quality control and data filtering pipelines used by Harris et al. as they applied the current state of the art quality thresholds to remove questionable sequences for detecting population level differences. Several mask files were applied to remove regions of the genome that might be lower quality, or might have very different mutation rates or base pair complexity compared to the rest of the genome. The 1kGP strict mask was used to remove low quality regions of the genome, highly conserved regions were removed using the phastCons100way mask file and highly repetitive regions were removed using the NestedRepeats mask file from RepeatMasker. Furthermore, only sites with missingness below 0.01, MAF less than 0.1, and MAF greater than 0.9 were considered. In total, 7,786,023 diallelic autosomal variants passed our quality controls for the mutation spectrum analysis. We calculated the mutation spectrum of base pair triplets for the list of significant variants for the JPT population using a similar method as described in (***Harris and Pritchard, 2017***).

For the reverse GWAS, the only filtration used was the application of an minor and major allele frequency cutoff of 0.000599 (removing singletons, doubletons and tripletons) resulting in a total of S=28,516,063 variants included in the test. We also used the NestedRepeats mask file to flag variants inside repetitive regions as these were analyzed separately for false discovery rate estimation. Variants flagged by the 1kGP strict mask are included in the association test and included in the FDR adjustment. These variants are only removed after the FDR and excluded from downstream discussion of error patterns, since most population genetics analyses use the strict mask as a filter, and we expect to find problematic variants in filtered regions.

### Testing the association of quality to genotype

When conducting a statistical analysis of population genetics data, we must account for population structure. In a typical GWAS, we are interested in modelling the phenotype as a function of the genotype. Here we have the opposite situation, where the quantitative variable (*Q*) is used as an explanatory variable. So we consider models where the genotype *y* is a function of an expected frequency *π*_*si*_, based on population structure, and *Q*. The null model is

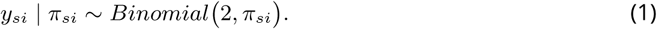

The expected frequency for a SNP *s* and individual *i* can be estimated using principal component analysis, categorical population labels, or logistic factor analysis (***Song et al., 2015***). The alternative model then takes in *Q* as a covariate:

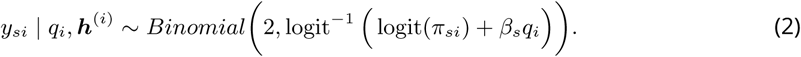

Under the null hypothesis the slope coefficient *β*_*s*_ is zero and Model (2) reduces to Model (1). *β*_*s*_ denotes the association to average quality of mapped bases *Q* to genotype *y*_*s*_. To test the null hypothesis, we use the generalized likelihood ratio test statistic, whose deviance is a measure of the marginal importance of adding *Q* in the model. The deviance test statistic under the null model is approximately chi-square distributed with one degrees of freedom.

We run a total of *S* regressions, where *S* is the total number of genomic loci. Given the large number of tests, the large proportion of expected null hypotheses and the positive dependencies across the genome, we used the two-stage Benjamini & Hochberg step-up FDR-controlling procedure to adjust the *p*-values (***Benjamini et al., 2006***). By using a nominal Type-I error rate *α* = 0.01, a total of 15,270 variants were found to be statistically significance. See Supplementary Data for a list of variants and adjusted *p*-values.

### Individual-specific allele frequency

Examples of models that are widely used to account population structure include the Balding-Nichols model (***Balding and Nichols, 1995***), and the Pritchard-Stephens-Donnelly model (***Pritchard et al., 2000***). These and several other similar models used in GWAS studies can be understood in terms of the following matrix factorization.

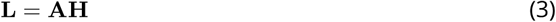

where the *i*^th^ column (***h***^(*i*)^) of the *K × I* matrix **H** encodes the population structure of the *i*^th^ individual and the *s*^th^ row of the *S × K* matrix **A** determines how that structure is manifested in SNP *s*. When Hardy-Weinberg equilibrium holds, observed genotype can be assumed to be generated by the following Binomial model.

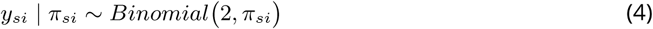

for *s* = 1 *…S* and *i* = *i, …, I*, where *y*_*si*_ ∈ {0, 1, 2} and *logit*(*π*_*si*_) is the (*s, i*) element of the matrix **L** such that *π*_*si*_ is the individual-specific allele frequency.

To test whether quality is associated to genotype while adjusting for population structure, we performed the Genotype-Conditional Association Test (GCAT) proposed by (***Song et al., 2015***). The GCAT is a regression approach that assumes the following model.

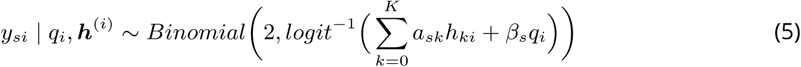

for *s* = 1 *…S* and *i* = *i, …, I* (*S* = 28, 516, 063 and *I* = 2, 504) and where 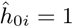 so that *a*_*s*0_ is the intercept term and 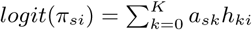. The vectors ***h***^*i*^ of the matrix **H** are unobserved but can be estimated using Logistic Factor Analysis (LFA) (***Song et al., 2015***) and are therefore used directly in the model. We approximated the population structure using *K* =5 latent components from a subsampled genotype matrix consisting of *M* = 2, 306, 130 SNPs (we picked SNPs from the 1kGP OMNI 2.5). To avoid possible biases in computing PCA from the biased variants, we considered the genotype matrix *L* obtained by downsampling 1kGP variants the positions from the OMNI 2.5M chip.

### Imputation

Using the Michigan Imputation Server, we imputed the genotype data from 1kGP for chromosomes 1 and 2. We used the genotyped data from the 1kGP Omni 2.5M chip genotype data. The VCF file returned from the server was then downloaded and used to search for the number of significant variants successfully imputed.

## Acknowledgments

We would like to thank K. Harris for sharing her mutation spectrum scripts, F. Mafessoni, G. Santpere and A. Navarro for feedback on earlier versions of the manuscript. We would also like to thank the members of the Gravel lab for their help with coding and useful discussions.

## References

1000 Genomes Project Consortium (2010). A map of human genome variation from population-scale sequencing. Nature, 467(7319):1061–73.

1000 Genomes Project Consortium (2012). An integrated map of genetic variation. Nature, 135:0–9.

Aikens, R. C., Johnson, K. E., and Voight, B. F. (2019). Signals of Variation in Human Mutation Rate at Multiple Levels of Sequence Context. Molecular Biology and Evolution.

Alexandrov, L. B., Nik-Zainal, S., Wedge, D. C., Aparicio, S. A., Behjati, S., Biankin, A. V., Bignell, G. R., Bolli, N., Borg, A., Børresen-Dale, A. L., Boyault, S., Burkhardt, B., Butler, A. P., Caldas, C., Davies, H. R., Desmedt, C., Eils, R., Eyfjörd, J. E., Foekens, J. A., Greaves, M., Hosoda, F., Hutter, B., Ilicic, T., Imbeaud, S., Imielinsk, M., Jäger, N., Jones, D. T., Jonas, D., Knappskog, S., Koo, M., Lakhani, S. R., López-Otín, C., Martin, S., Munshi, N. C., Nakamura, H., Northcott, P. A., Pajic, M., Papaemmanuil, E., Paradiso, A., Pearson, J. V., Puente, X. S., Raine, K., Ramakrishna, M., Richardson, A. L., Richter, J., Rosenstiel, P., Schlesner, M., Schumacher, T. N., Span, P. N., Teague, J. W., Totoki, Y., Tutt, A. N., Valdés-Mas, R., Van Buuren, M. M., Van ’T Veer, L., Vincent-Salomon, A., Waddell, N., Yates, L. R., Zucman-Rossi, J., Andrew Futreal, P., McDermott, U., Lichter, P., Meyerson, M., Grimmond, S. M., Siebert, R., Campo, E., Shibata, T., Pfister, S. M., Campbell, P. J., and Stratton, M. R. (2013). Signatures of mutational processes in human cancer. Nature, 500(7463):415–421.

Astle, W. J., Elding, H., Jiang, T., Allen, D., Ruklisa, D., Mann, A. L., Mead, D., Bouman, H., Riveros-Mckay, F., Kostadima, M. A., et al. (2016). The allelic landscape of human blood cell trait variation and links to common complex disease. Cell, 167(5):1415–1429.

Balding, D. J. and Nichols, R. A. (1995). A method for quantifying differentiation between populations at multi-allelic loci and its implications for investigating identity and paternity. Genetica, 96(1-2):3–12.

Benjamini, Y., Krieger, A. M., and Yekutieli, D. (2006). Adaptive linear step-up procedures that control the false discovery rate. Biometrika.

Consortium, . G. P. et al. (2015). A global reference for human genetic variation. Nature, 526(7571):68.

Ebejer, J. L., Duffy, D. L., Van Der Werf, J., Wright, M. J., Montgomery, G., Gillespie, N. A., Hickie, I. B., Martin, N. G., and Medland, S. E. (2013). Genome-wide association study of inattention and hyperactivity-impulsivity measured as quantitative traits. Twin Research and Human Genetics.

Ellinghaus, D., Jostins, L., Spain, S. L., Cortes, A., Bethune, J., Han, B., Park, Y. R., Raychaudhuri, S., Pouget, J. G., Hübenthal, M., et al. (2016). Analysis of 1ve chronic in2ammatory diseases identi1es 27 new associations and highlights disease-specific patterns at shared loci. Nature genetics, 48(5):510.

Gao, X. R., Huang, H., Nannini, D. R., Fan, F., and Kim, H. (2018). Genome-wide association analyses identify new loci influencing intraocular pressure. Human molecular genetics, 27(12):2205–2213.

Harris, K. (2015). Evidence for recent, population-specific evolution of the human mutation rate. Proceedings of the National Academy of Sciences, 112(11):3439–3444.

Harris, K. and Pritchard, J. K. (2017). Rapid evolution of the human mutation spectrum. eLife, 6.

Herold, C., Hooli, B. V., Mullin, K., Liu, T., Roehr, J. T., Mattheisen, M., Parrado, A. R., Bertram, L., Lange, C., and Tanzi, R. E. (2016). Family-based association analyses of imputed genotypes reveal genome-wide significant association of alzheimer?s disease with osbpl6, ptprg, and pdcl3. Molecular psychiatry, 21(11):1608.

International HapMap Consortium (2005). A haplotype map of the human genome. Nature, 437(7063):1299–320.

Kraja, A. T., Vaidya, D., Pankow, J. S., Goodarzi, M. O., Assimes, T. L., Kullo, I. J., Sovio, U., Mathias, R. A., Sun, Y. V., Franceschini, N., Absher, D., Li, G., Zhang, Q., Feitosa, M. F., Glazer, N. L., Haritunians, T., Hartikainen, A. L., Knowles, J. W., North, K. E., Iribarren, C., Kral, B., Yanek, L., O’Reilly, P. F., McCarthy, M. I., Jaquish, C., Couper, D. J., Chakravarti, A., Psaty, B. M., Becker, L. C., Province, M. A., Boerwinkle, E., Quertermous, T., Palotie, L., Jarvelin, M. R., Becker, D. M., Kardia, S. L., Rotter, J. I., Chen, Y. D. I., and Borecki, I. B. (2011). A bivariate genome-wide approach to metabolic syndrome: STAMPEED Consortium. Diabetes.

Lan, T., Lin, H., Zhu, W., Laurent, T. C. A. M., Yang, M., Liu, X., Wang, J., Wang, J., Yang, H., Xu, X., and Guo, X. (2017). Deep whole-genome sequencing of 90 han chinese genomes. GigaScience, 6(9):gix067.

Lek, M., Karczewski, K. J., Minikel, E. V., Samocha, K. E., Banks, E., Fennell, T., O’Donnell-Luria, A. H., Ware, J. S., Hill, A. J., Cummings, B. B., Tukiainen, T., Birnbaum, D. P., Kosmicki, J. A., Duncan, L. E., Estrada, K., Zhao, F., Zou, J., Pierce-Hoffman, E., Berghout, J., Cooper, D. N., Deflaux, N., DePristo, M., Do, R., Flannick, J., Fromer, M., Gauthier, L., Goldstein, J., Gupta, N., Howrigan, D., Kiezun, A., Kurki, M. I., Moonshine, A. L., Natarajan, P., Orozco, L., Peloso, G. M., Poplin, R., Rivas, M. A., Ruano-Rubio, V., Rose, S. A., Ruderfer, D. M., Shakir, K., Stenson, P. D., Stevens, C., Thomas, B. P., Tiao, G., Tusie-Luna, M. T., Weisburd, B., Won, H. H., Yu, D., Altshuler, D. M., Ardissino, D., Boehnke, M., Danesh, J., Donnelly, S., Elosua, R., Florez, J. C., Gabriel, S. B., Getz, G., Glatt, S. J., Hultman, C. M., Kathiresan, S., Laakso, M., McCarroll, S., McCarthy, M. I., McGovern, D., McPherson, R., Neale, B. M., Palotie, A., Purcell, S. M., Saleheen, D., Scharf, J. M., Sklar, P., Sullivan, P. F., Tuomilehto, J., Tsuang, M. T., Watkins, H. C., Wilson, J. G., Daly, M. J., and MacArthur, D. G. (2016). Analysis of protein-coding genetic variation in 60,706 humans. Nature, 536(7616):285–291.

López-Mejías, R., Carmona, F. D., Castañeda, S., Genre, F., Remuzgo-Martínez, S., Sevilla-Perez, B., Ortego-Centeno, N., Llorca, J., Ubilla, B., Mijares, V., et al. (2017). A genome-wide association study suggests the hla class ii region as the major susceptibility locus for iga vasculitis. Scientific reports, 7(1):5088.

Lutz, S. M., Cho, M. H., Young, K., Hersh, C. P., Castaldi, P. J., McDonald, M.-L., Regan, E., Mattheisen, M., DeMeo, D. L., Parker, M., et al. (2015). A genome-wide association study identifies risk loci for spirometric measures among smokers of european and african ancestry. BMC genetics, 16(1):138.

Mafessoni, F., Prasad, R. B., Groop, L., Hansson, O., and Prüfer, K. (2018). Turning vice into virtue: Using batch-effects to detect errors in large genomic data sets. Genome biology and evolution, 10(10):2697–2708.

Mallick, S., Li, H., Lipson, M., Mathieson, I., Gymrek, M., Racimo, F., Zhao, M., Chennagiri, N., Nordenfelt, S., Tandon, A., Skoglund, P., Lazaridis, I., Sankararaman, S., Fu, Q., Rohland, N., Renaud, G., Erlich, Y., Willems, T., Gallo, C., Spence, J. P., Song, Y. S., Poletti, G., Balloux, F., Van Driem, G., De Knijff, P., Romero, I. G., Jha, A. R., Behar, D. M., Bravi, C. M., Capelli, C., Hervig, T., Moreno-Estrada, A., Posukh, O. L., Balanovska, E., Balanovsky, O., Karachanak-Yankova, S., Sahakyan, H., Toncheva, D., Yepiskoposyan, L., Tyler-Smith, C., Xue, Y., Abdullah, M. S., Ruiz-Linares, A., Beall, C. M., Di Rienzo, A., Jeong, C., Starikovskaya, E. B., Metspalu, E., Parik, J., Villems, R., Henn, B. M., Hodoglugil, U., Mahley, R., Sajantila, A., Stamatoyannopoulos, G., Wee, J. T., Khusainova, R., Khusnutdinova, E., Litvinov, S., Ayodo, G., Comas, D., Hammer, M. F., Kivisild, T., Klitz, W., Winkler, C. A., Labuda, D., Bamshad, M., Jorde, L. B., Tishkoff, S. A., Watkins, W. S., Metspalu, M., Dryomov, S., Sukernik, R., Singh, L., Thangaraj, K., Paäbo, S., Kelso, J., Patterson, N., and Reich, D. (2016). The Simons Genome Diversity Project: 300 genomes from 142 diverse populations. Nature, 538(7624):201–206.

Mandage, R., Telford, M., Rodríguez, J. A., Farré, X., Layouni, H., Marigorta, U. M., Cundiff, C., Heredia-Genestar, J. M., Navarro, A., and Santpere, G. (2017). Genetic factors affecting EBV copy number in lymphoblastoid cell lines derived from the 1000 Genome Project samples. PLoS ONE.

Mathieson, I. and Reich, D. (2017). Differences in the rare variant spectrum among human populations. PLoS Genetics, 13(2).

McCarthy, S., Das, S., Kretzschmar, W., Delaneau, O., Wood, A. R., Teumer, A., Kang, H. M., Fuchsberger, C., Danecek, P., Sharp, K., et al. (2016). A reference panel of 64,976 haplotypes for genotype imputation. Nature genetics, 48(10):1279.

Minoche, A. E., Dohm, J. C., and Himmelbauer, H. (2011). Evaluation of genomic high-throughput sequencing data generated on Illumina HiSeq and Genome Analyzer systems. Genome Biology, 12(11).

Nagy, R., Boutin, T. S., Marten, J., Huffman, J. E., Kerr, S. M., Campbell, A., Evenden, L., Gibson, J., Amador, C., Howard, D. M., et al. (2017). Exploration of haplotype research consortium imputation for genome-wide association studies in 20,032 generation scotland participants. Genome medicine, 9(1):23.

Nishida, N., Sugiyama, M., Sawai, H., Nishina, S., Sakai, A., Ohashi, J., Khor, S.-S., Kakisaka, K., Tsuchiura, T., Hino, K., et al. (2018). Key hla-drb1-dqb1 haplotypes and role of the btnl2 gene for response to a hepatitis b vaccine. Hepatology, 68(3):848–858.

Park, S. L., Carmella, S. G., Chen, M., Patel, Y., Stram, D. O., Haiman, C. A., Le Marchand, L., and Hecht, S. S. (2015). Mercapturic acids derived from the toxicants acrolein and crotonaldehyde in the urine of cigarette smokers from five ethnic groups with differing risks for lung cancer. PLoS One, 10(6):e0124841.

Pfeifer, G. P., Denissenko, M. F., Olivier, M., Tretyakova, N., Hecht, S. S., and Hainaut, P. (2002). Tobacco smoke carcinogens, DNA damage and p53 mutations in smoking-associated cancers. Oncogene, 21-48(6):7435–7451.

Pleasance, E. D., Stephens, P. J., O’Meara, S., McBride, D. J., Meynert, A., Jones, D., Lin, M. L., Beare, D., Lau, K. W., Greenman, C., Varela, I., Nik-Zainal, S., Davies, H. R., Ordõez, G. R., Mudie, L. J., Latimer, C., Edkins, S., Stebbings, L., Chen, L., Jia, M., Leroy, C., Marshall, J., Menzies, A., Butler, A., Teague, J. W., Mangion, J., Sun, Y. A., McLaughlin, S. F., Peckham, H. E., Tsung, E. F., Costa, G. L., Lee, C. C., Minna, J. D., Gazdar, A., Birney, E., Rhodes, M. D., McKernan, K. J., Stratton, M. R., Futreal, P. A., and Campbell, P. J. (2010). A small-cell lung cancer genome with complex signatures of tobacco exposure. Nature, 463(7278):184–190.

Pritchard, J. K., Stephens, M., and Donnelly, P. (2000). Inference of population structure using multilocus genotype data. Genetics, 155(2):945–959.

Shiraishi, Y., Tremmel, G., Miyano, S., and Stephens, M. (2015). A Simple Model-Based Approach to Inferring and Visualizing Cancer Mutation Signatures. PLoS Genetics, 11(12).

Song, M., Hao, W., and Storey, J. D. (2015). Testing for genetic associations in arbitrarily structured populations. Nature genetics, 47(5):550.

Spracklen, C. N., Chen, P., Kim, Y. J., Wang, X., Cai, H., Li, S., Long, J., Wu, Y., Wang, Y. X., Takeuchi, F., et al. (2017). Association analyses of east asian individuals and trans-ancestry analyses with european individuals reveal new loci associated with cholesterol and triglyceride levels. Human molecular genetics, 26(9):1770–1784.

Suhre, K., Arnold, M., Bhagwat, A. M., Cotton, R. J., Engelke, R., Raffler, J., Sarwath, H., Thareja, G., Wahl, A., DeLisle, R. K., et al. (2017). Connecting genetic risk to disease end points through the human blood plasma proteome. Nature communications, 8:14357.

Tian, C., Hromatka, B. S., Kiefer, A. K., Eriksson, N., Noble, S. M., Tung, J. Y., and Hinds, D. A. (2017). Genome-wide association and hla region fine-mapping studies identify susceptibility loci for multiple common infections. Nature communications, 8(1):599.

van Dijk, E. L., Auger, H., Jaszczyszyn, Y., and Thermes, C. (2014). Ten years of next-generation sequencing technology. Trends in Genetics, 30(9):418–426.

Xu, J., Mo, Z., Ye, D., Wang, M., Liu, F., Jin, G., Xu, C., Wang, X., Shao, Q., Chen, Z., et al. (2012). Genome-wide association study in chinese men identifies two new prostate cancer risk loci at 9q31. 2 and 19q13. 4. Nature genetics, 44(11):1231.

Yucesoy, B., Kaufman, K. M., Lummus, Z. L., Weirauch, M. T., Zhang, G., Cartier, A., Boulet, L.-P., Sastre, J., Quirce, S., Tarlo, S. M., et al. (2015). Genome-wide association study identifies novel loci associated with diisocyanate-induced occupational asthma. Toxicological Sciences, 146(1):192–201.

