## Supplementary Figures for "Legacy Data Confounds Genomics Studies"

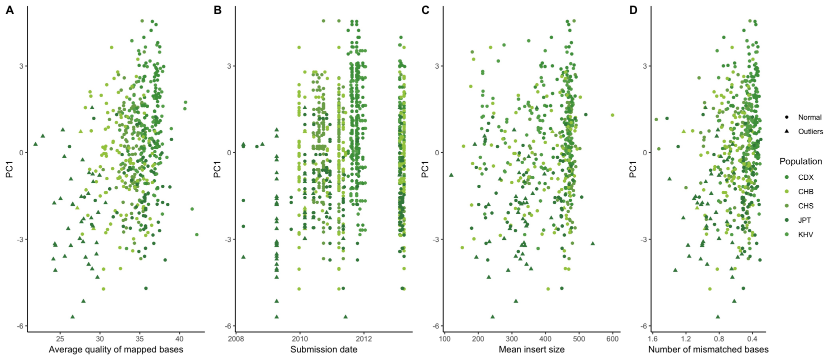


**Figure S1.** Sequencing metrics against the prevalence of the *AC→*CC mutational signature in 1000 Genomes Project. The average quality per mapped bases $Q$ per individual shows some clustering with individuals with low-quality data showing elevated rates of the signature.

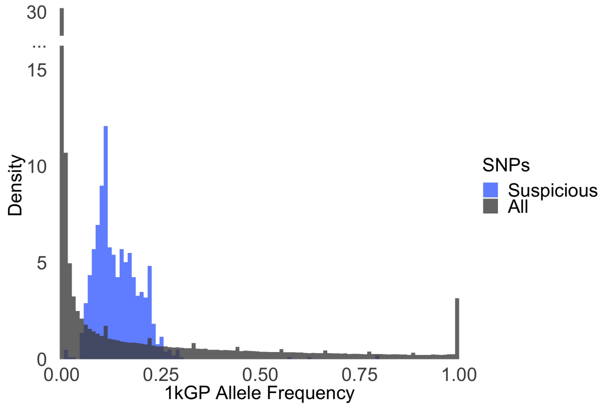


**Figure S2.** Frequency distribution of the 1034 $Q$ associated SNPs from the single population test in the JPT in blue and all SNPs in black. The association test lacks power to identify variants at low frequencies.

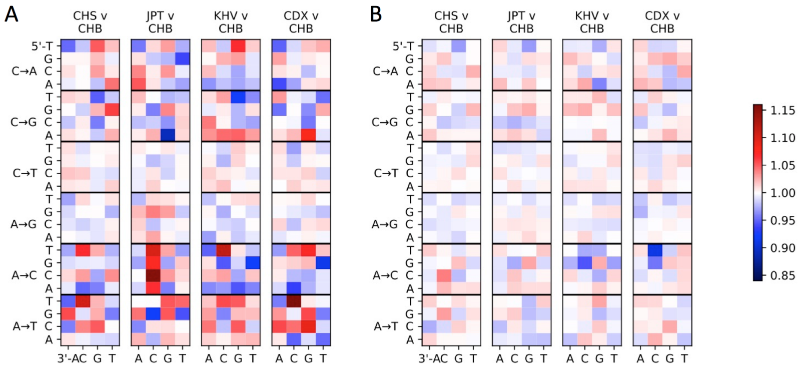


**Figure S3.** Comparing mutational signatures after removing $Q$-associated variants and after removing individuals with low $Q$. **A** The *AC→*CC mutational signature in JPT remains despite removing variants associated to quality. **B** Removing individuals with average quality per mapped bases $Q$ below a threshold of 30 removes the mutational signature completely.

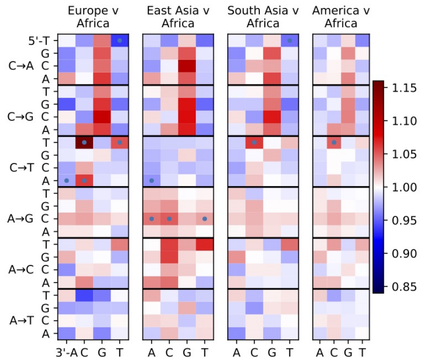


**Figure S4.** Comparing continental mutational signatures after removing $Q$-associated variants and after removing individuals with low $Q$. The *AC→*CC mutational signature in East Asian (EAS) Population remains despite removing individuals with average quality per mapped bases $Q$ below a threshold of 30. Points indicate significant mutation spectrum contrasts with $p < 10^{-5}$

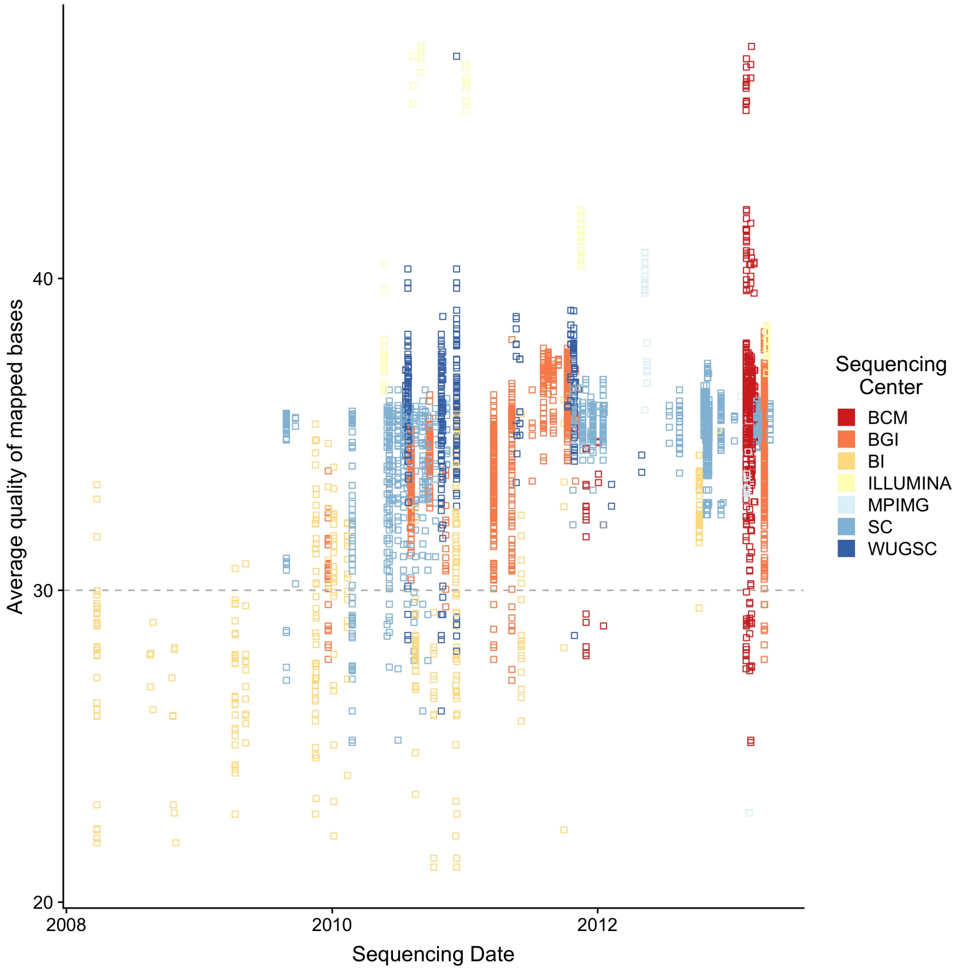


**Figure S5.** Sequencing date of samples included in the 1kGP coloured by sequencing centre. Improvements in sequencing platforms coincides with improvement in sequencing quality.

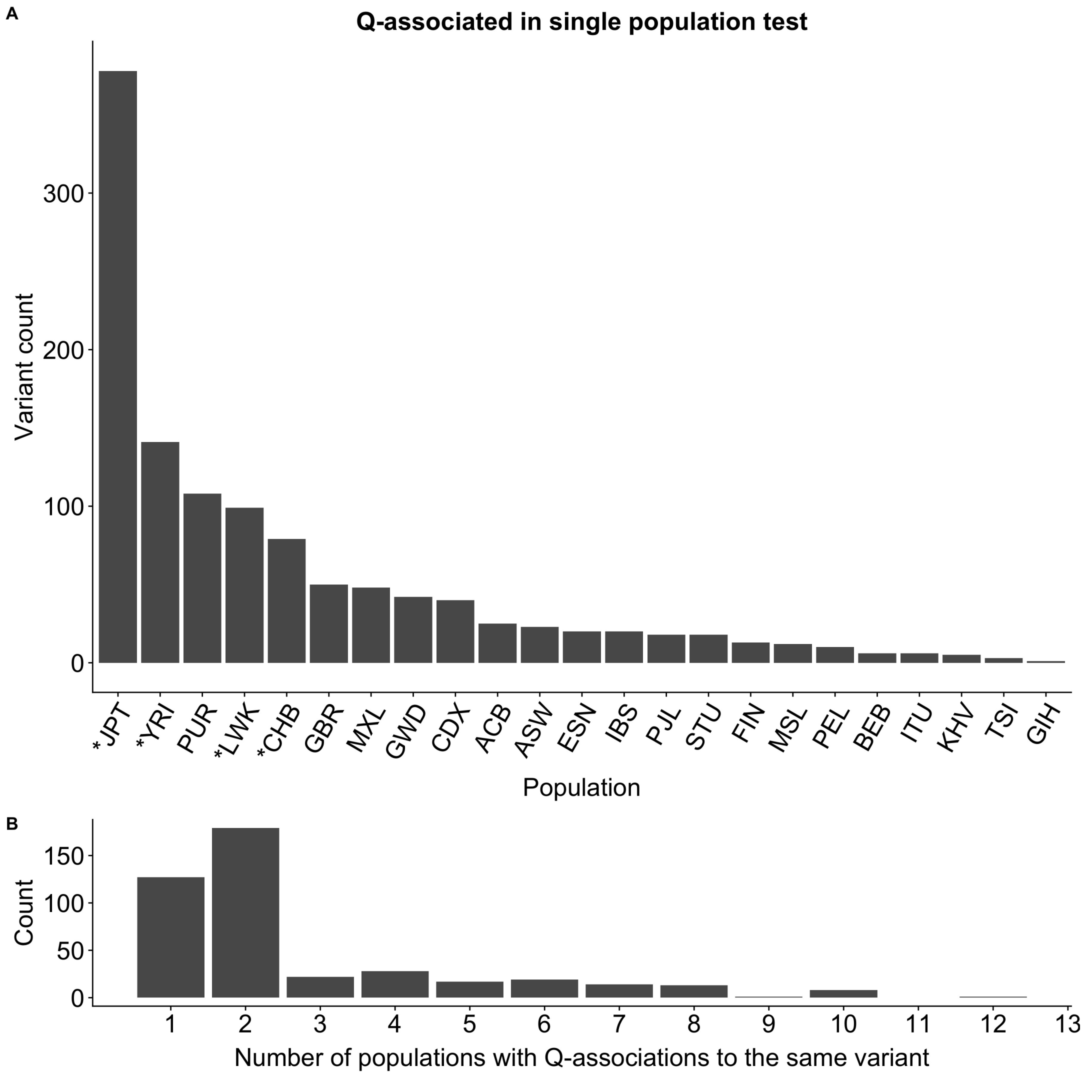


**Figure S6.** Results from the single population test. We find a total of 1,165 variants associated with the average quality of mapped bases $Q$. Phase 1 populations are indicated by a star ( * ). **A** For each population, we plot the number of variants associated to $Q$. **B** The number of populations for which a variant has reached significance.

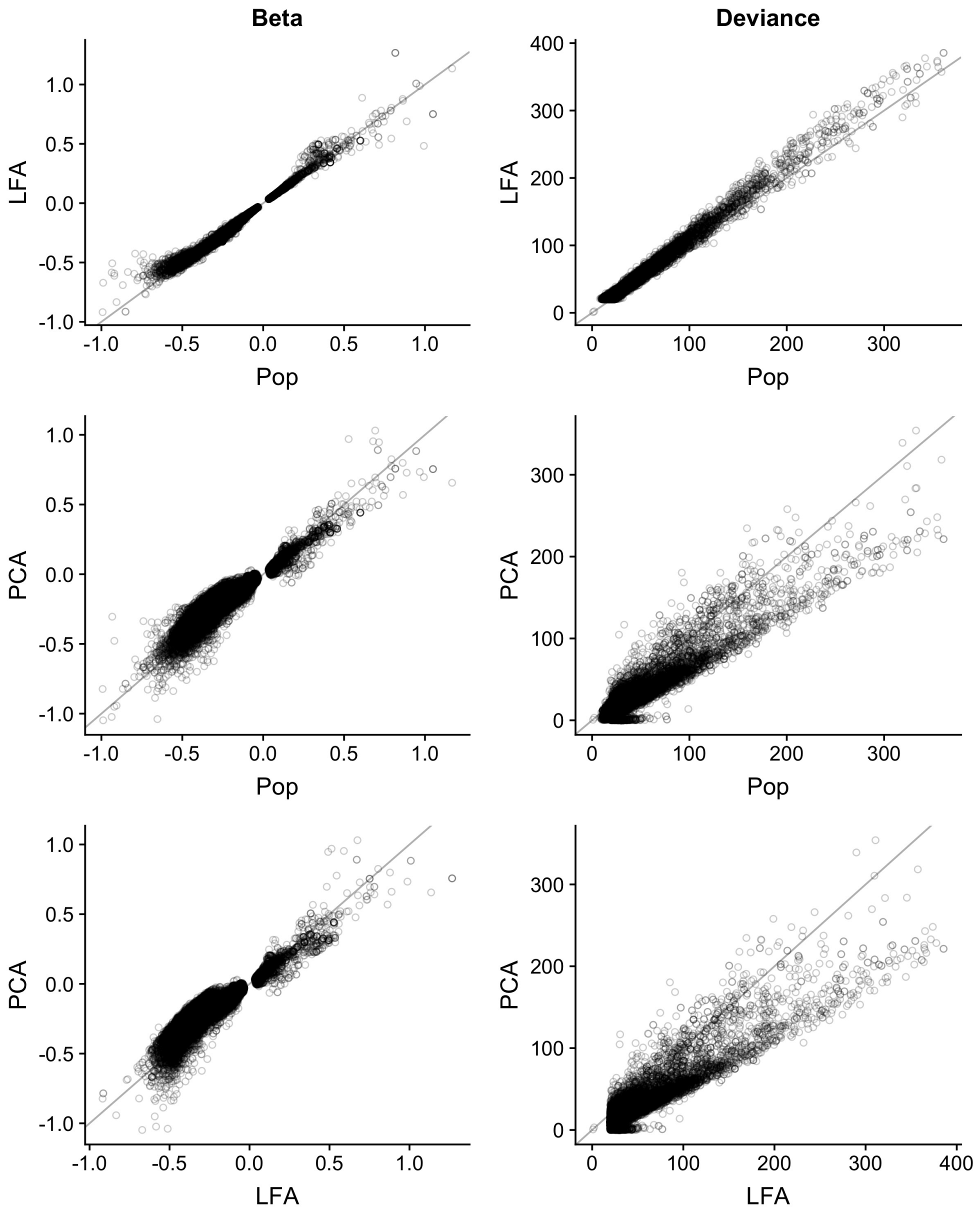


**Figure S7.** Comparison of three logistic regression models for testing association to $Q$. These methods model each genotype as a logistic function using principal components (PC), Population membership (Pop) or LFA as an offset. In these plots we are comparing the deviance from the null model in the 15,270 variants identified using the LFA model.

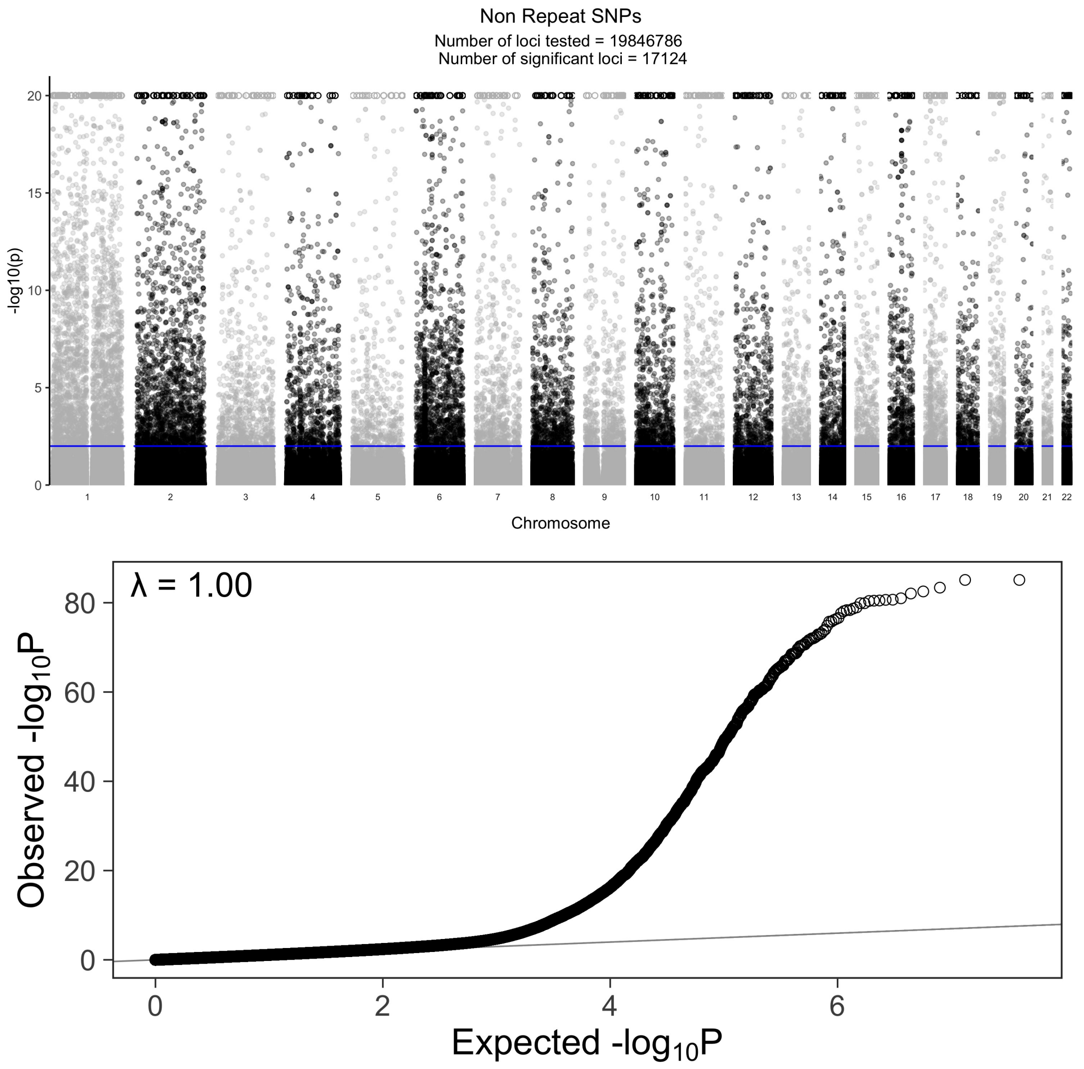


**Figure S8.** Association of SNPS in non-repetitive regions with $Q$. **A** Manhattan plot of the $-\log_{10}(p)$ values for the reverse GWAS logistic regression analysis for SNPs in non repetitive regions. There are 15,018 SNPs that reach $p$ values greater than $p < 0.01$ after performing a two-stage Benjamini and Hochberg FDR adjustment. The circles ( o ) are variants that reached values greater than 20, for clarity we implemented hard ceiling at 20. **B** QQ plot of the unadjusted p values for the reverse GWAS logistic regression analysis for SNPs in non repetitive regions.

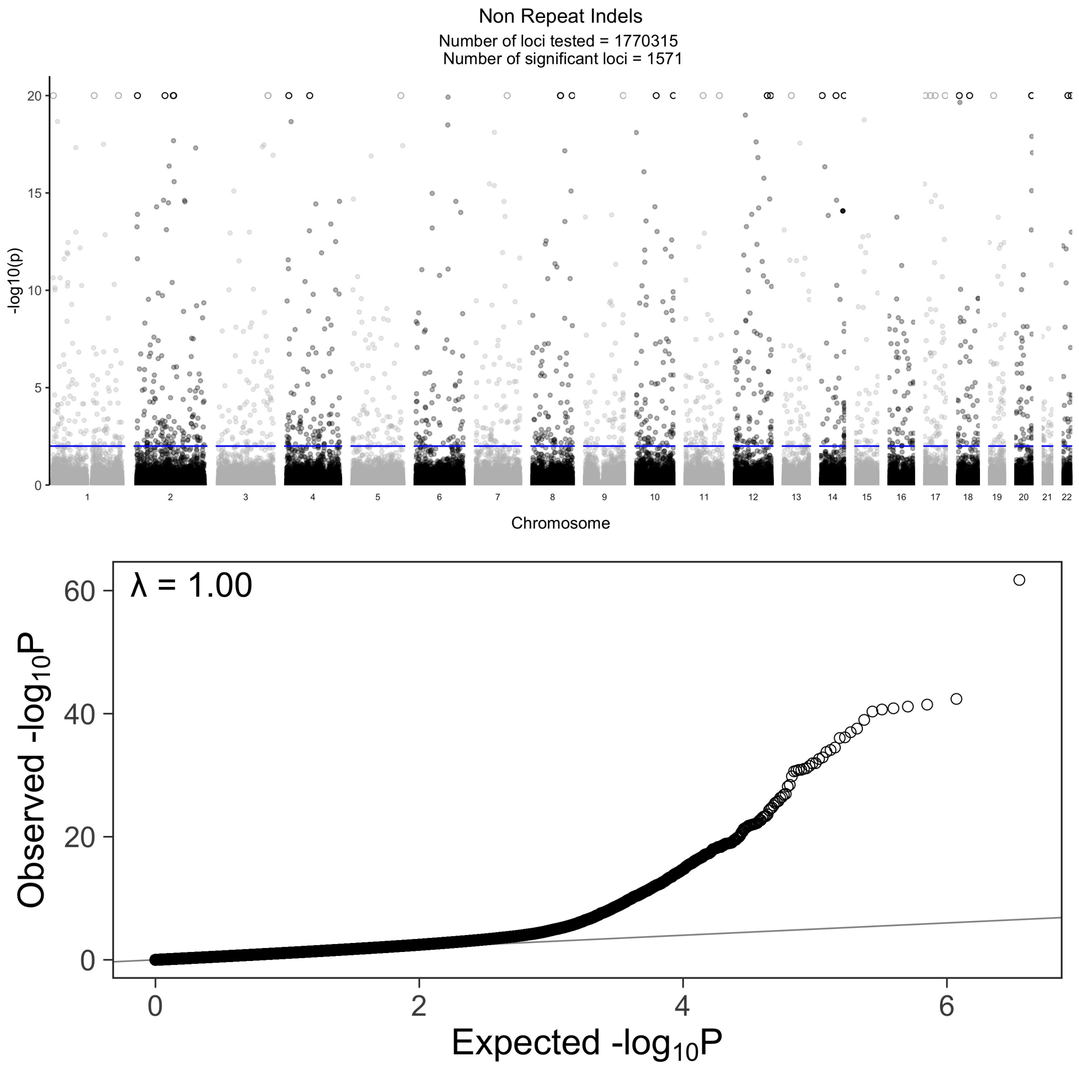


**Figure S9.** Association of indels in non-repetitive regions with $Q$. **A** Manhattan plot of the $-\log_{10}(p)$ values for the reverse GWAS logistic regression analysis for INDELs in non repetitive regions. There are 2,121 INDELs that reach $p$ values greater than $p < 0.01$ after performing a two-stage Benjamini and Hochberg FDR adjustment. The circles ( o ) are variants that reached values greater than 20, for clarity we implemented hard ceiling at 20. **B** QQ plot of the unadjusted p values for the reverse GWAS logistic regression analysis for INDELs in non repetitive regions.

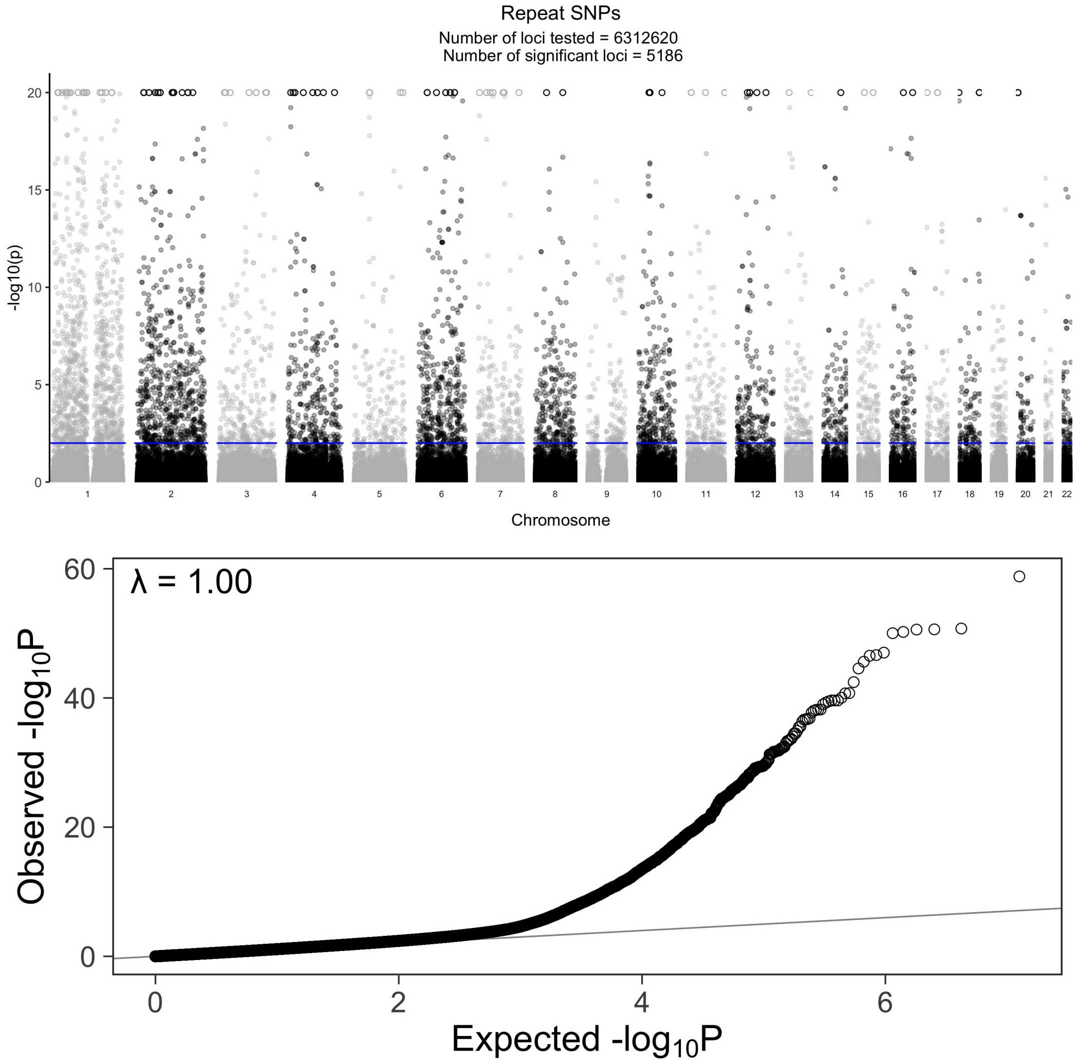


**Figure S10.** Association of SNPS in repetitive regions with $Q$. **A** Manhattan plot of the $-\log_{10}(p)$ values for the reverse GWAS logistic regression analysis for SNPs in repetitive regions. There are 4,405 SNPs that reach $p$ values greater than $p < 0.01$ after performing a two-stage Benjamini and Hochberg FDR adjustment. The circles ( o ) are variants that reached values greater than 20, for clarity we implemented hard ceiling at 20. **B** QQ plot of the unadjusted p values for the reverse GWAS logistic regression analysis for SNPs in repetitive regions.

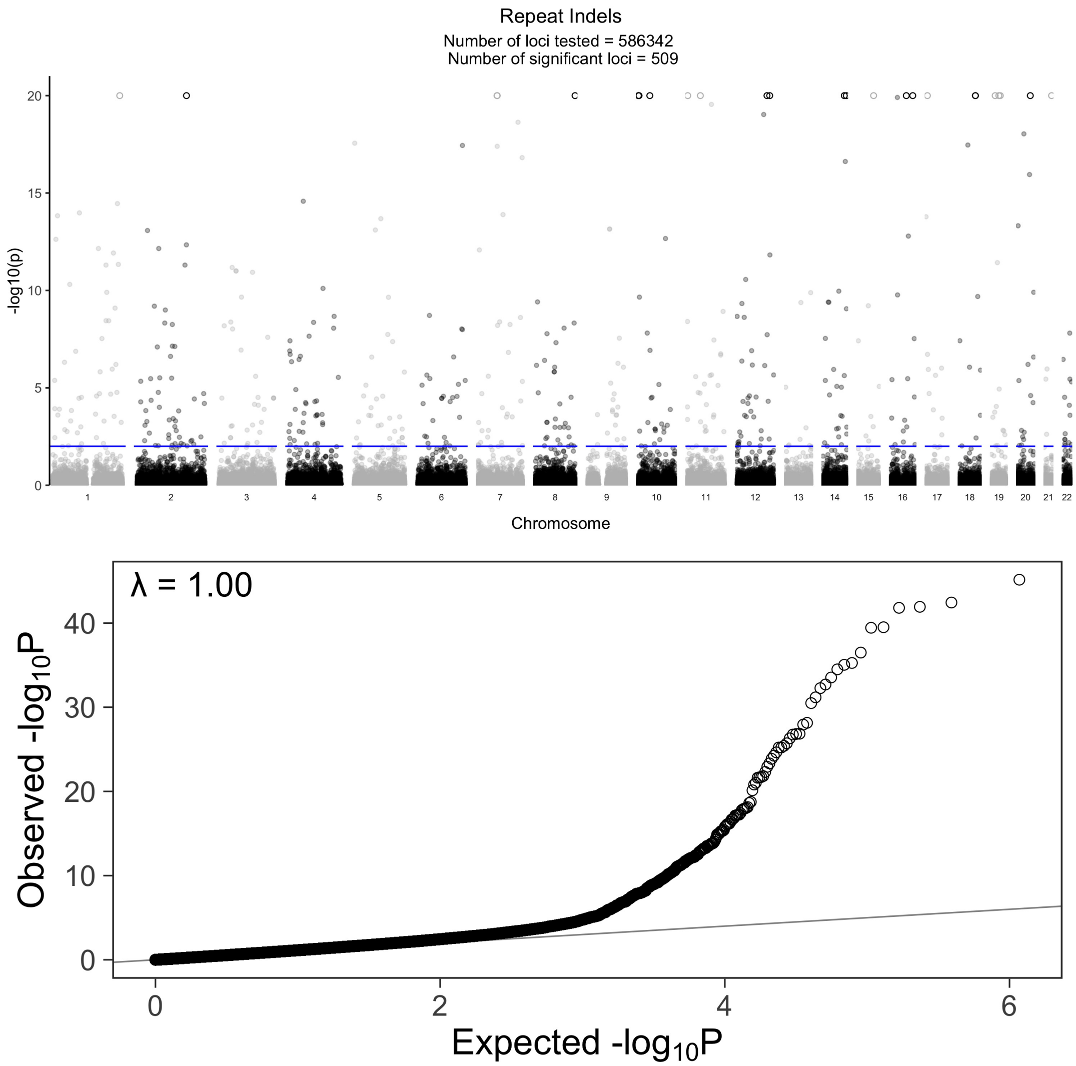


**Figure S11.** Association of indels in repetitive regions with $Q$. **A** Manhattan plot of the $-\log_{10}(p)$ values for the reverse GWAS logistic regression analysis for INDELs in repetitive regions. There are 642 INDELs that reach $p$ values greater than $p < 0.01$ after performing a two-stage Benjamini and Hochberg FDR adjustment. The circles ( o ) are variants that reached values greater than 20, for clarity we implemented hard ceiling at 20. **B** QQ plot of the unadjusted p values for the reverse GWAS logistic regression analysis for INDELs in repetitive regions.

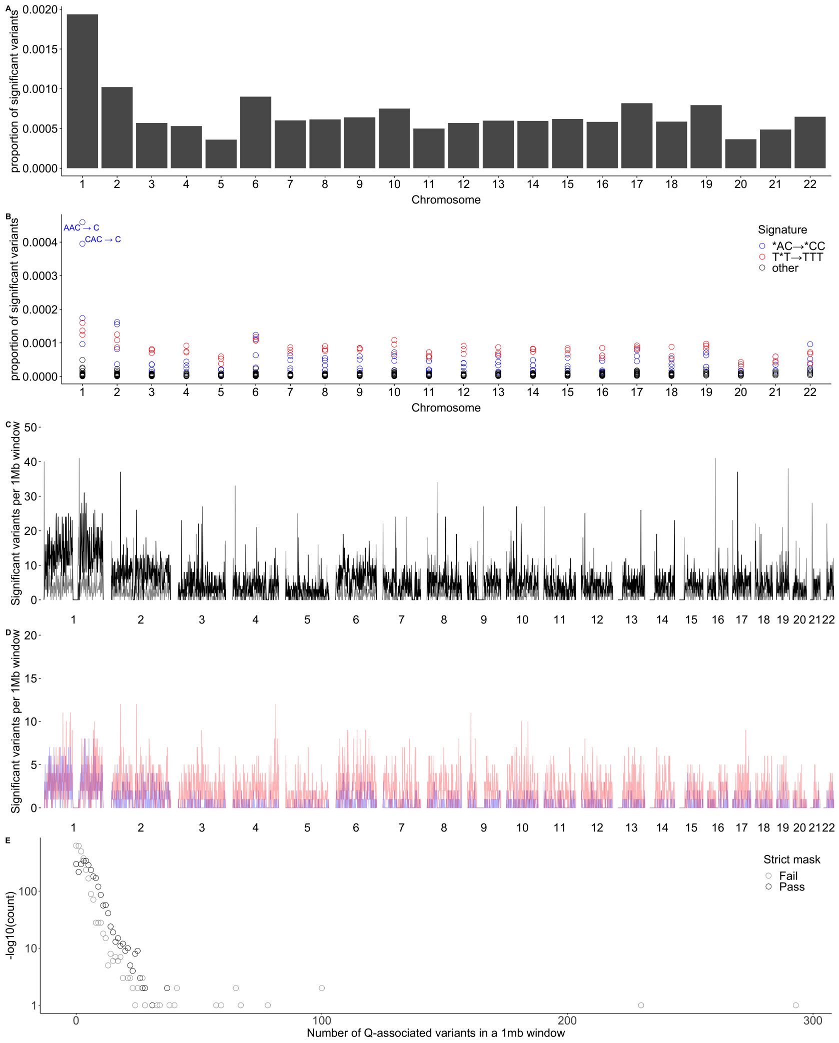


**Figure S12.** **A** The proportion of $Q$-associated variants per chromosome. **B** Mutation spectrum enrichment of $Q$-associated variants across each chromosome. Chromosome 1 has a strong enrichment in *AC→*CC mutational pattern. **C** The number of $Q$-associated variants per 1Mb window across the genome. Grey bars indicate regions within and black bars indicate regions outside the 1000 Genomes Project strict mask. **D** Blue bars indicate the density of variants passing the strict mask with the *AC→*CC mutational pattern whereas red points have the T*T→TTT mutational pattern. Three regions not flagged by the 1000 Genomes Project strict mask in chromosomes 1, 2 and 17 have more than 30 variants per window. **E** Distribution of variant counts per 1Mb window. Where variants that fail the strict mask are in grey and those that pass are in black.

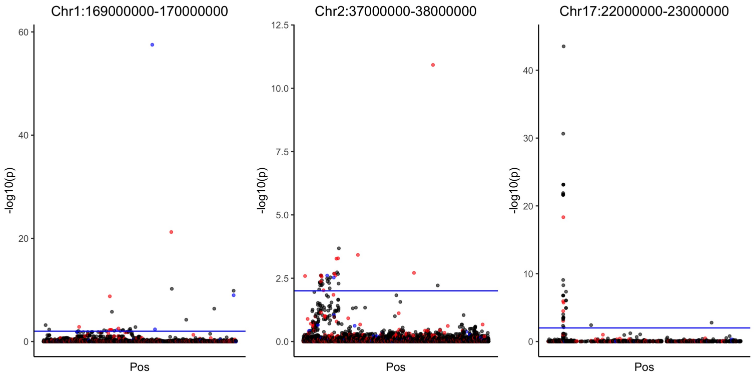


**Figure S13.** Manhattan plot of the $-\log_{10}(p)$ values for the reverse GWAS logistic regression analysis for the three 1Mb windows with the most $Q$-associated variants per 1Mb across the genome. In these plots, blue points are variants with the *AC→*CC mutational pattern whereas red points have the T*T→TTT mutational pattern.

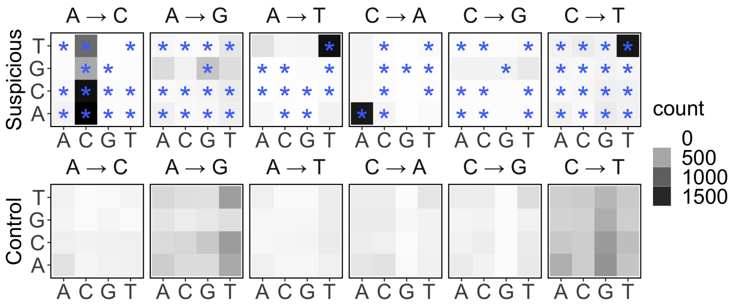


**Figure S14.** Mutation spectrum of the 15,270 $Q$-associated variants that pass the 1kGP strict mask filter, compared to a random set of non-associated variants. A large fraction of the $Q$-associated variants have the *AC→*CC mutational pattern. There is also an enrichment in TAT→TTT, TCT→TTT and TGT→TTT (shown as reverse complement ACA→AAA). These three enrichments can be summarized as T*T→TTT. Stars ( * ) indicate significant difference from the mutational spectrum of a set of control variants of equal size.

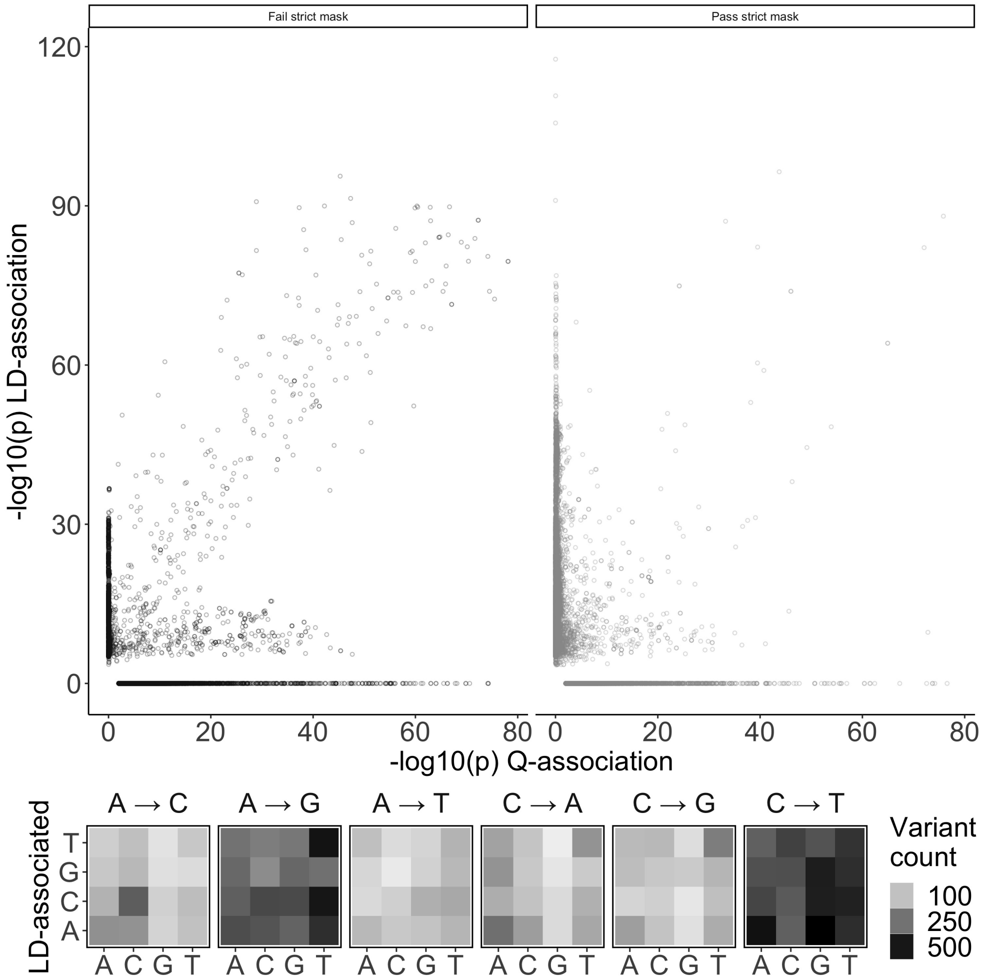


**Figure S15.** **A** Comparison of the p-values from the association to unusual LD from **?** and the p-values from the GCAT association test to quality metrics. There are a total of 42,307 variants in this plot with only 1,279 ( 3% ) are statistically significant in both tests. There are 23,111 variants ( 55% ) that are $Q$-associated but not LD-associated and 17,917 ( 42% ) that are LD-associated but not $Q$-associated. For the 21,416 variants that pass the strict mask, 629 ( 3% ) variants are both $Q$-associated and LD-associated; 8,495 variants ( 40% ) are $Q$-associated but not LD-associated and 12,292 ( 57% ) are LD-associated but not $Q$-associated. For the 14,616 variants that don't pass the strict mask, 650 ( 3% ) variants are both $Q$-associated and LD-associated; 14,616 variants ( 70% ) are $Q$-associated but not LD-associated and 5,625 ( 27% ) are LD-associated but not $Q$-associated. **B** Mutation spectrum of the variants significantly associated to unusual LD from **?**.

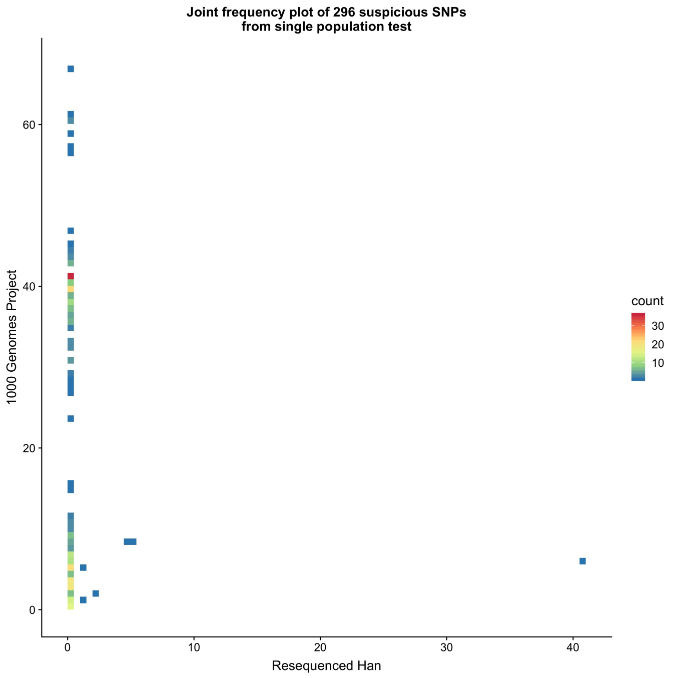


**Figure S16.** Site frequency spectrum plot comparing the original 1000 Genomes Project data to the high depth resequence data for variants that, in the 1000 Genomes Project, are both associated with $Q$ and polymorphic in the 83 individuals that were resequenced. Among the 296 variants associated with $Q$ in the single population tests within the 1000 Genomes Project CHB and CHS, 6 are present in the resequenced data (**?**).

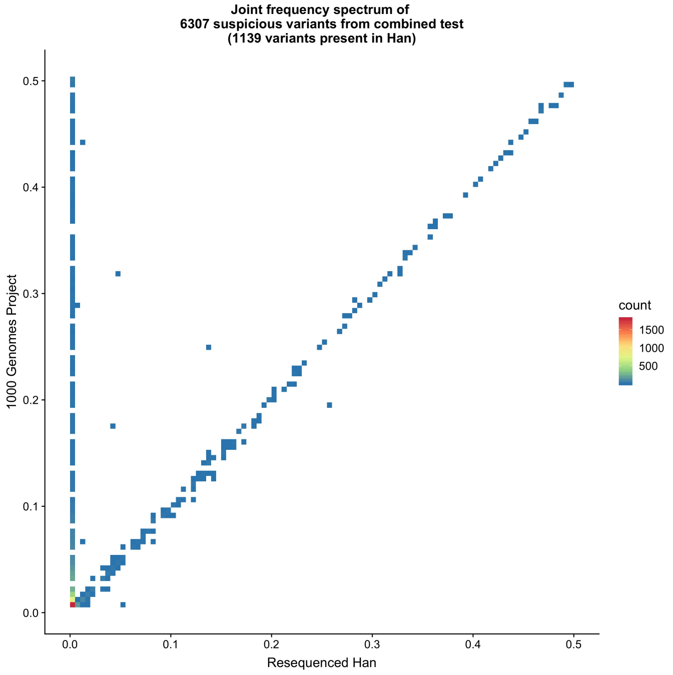


**Figure S17.** Site frequency spectrum plot comparing the original 1000 Genomes Project data to the high depth resequence data for variants that, in the 1000 Genomes Project, are both associated with $Q$ and polymorphic in the 83 individuals that were resequenced. Among the 6,307 variants associated with $Q$ in the GCAT model including all populations, 1,139 are present in the high depth resequenced individuals.

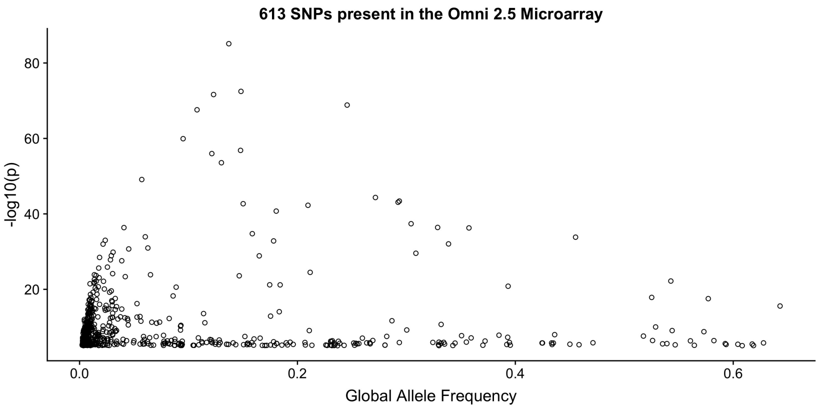


**Figure S18.** Estimated frequency and association strength of $Q$- associated variants present on Illumina's Omni 2.5 chip. Variants highly associated to $Q$ tend to have low global allele frequencies.

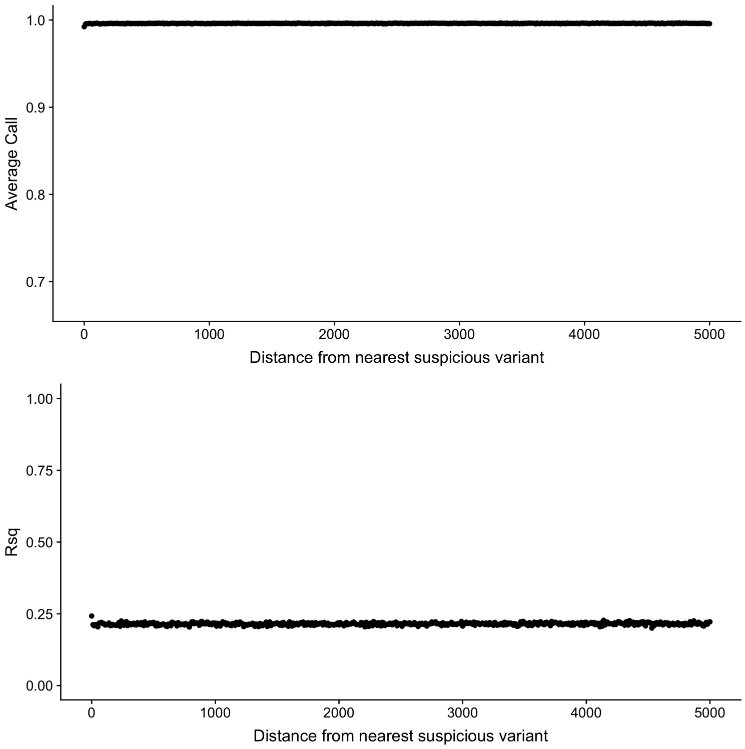


**Figure S19.** Imputation scores are not impacted by nearby suspicious variants. By comparing the imputation scores from Japanese Nagahama genotype data to the distance from the nearest suspicious variant, we find no impact on nearby variants. The only variants that do appear to have lower imputation scores are the suspicious variants themselves with a distance of zero on the plot.

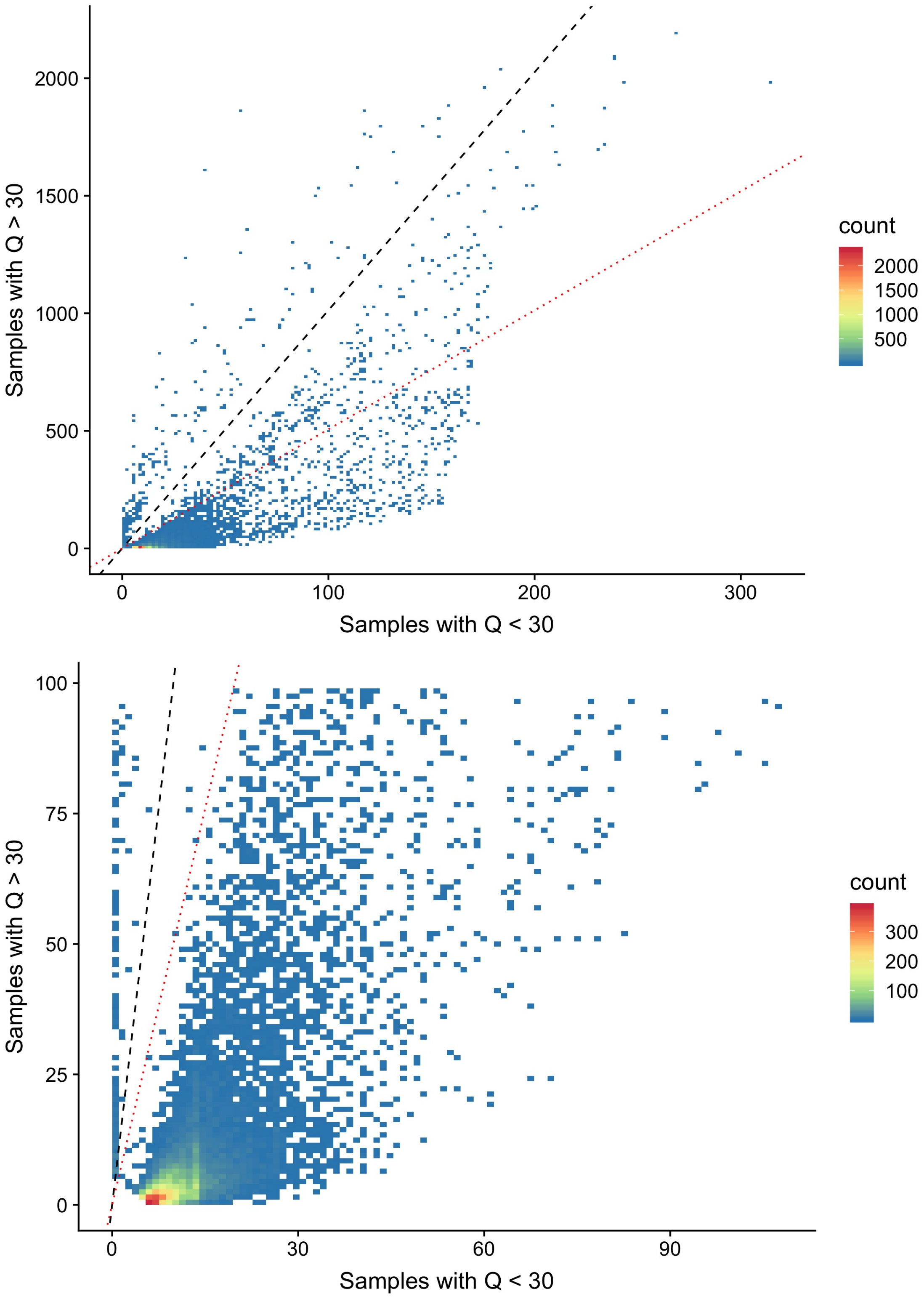


**Figure S20.** Site frequency spectrum plot comparing the allele frequency difference between individuals with low- and high-$Q$. The black dashed lines indicates equal allele frequencies while the red dotted line for variants twice as frequent in individuals with $Q$ scores below 30. Two clusters of are visible, where the majority (92.7%) of the $Q$-associated variants are more than twice as frequent in individuals with low-$Q$.

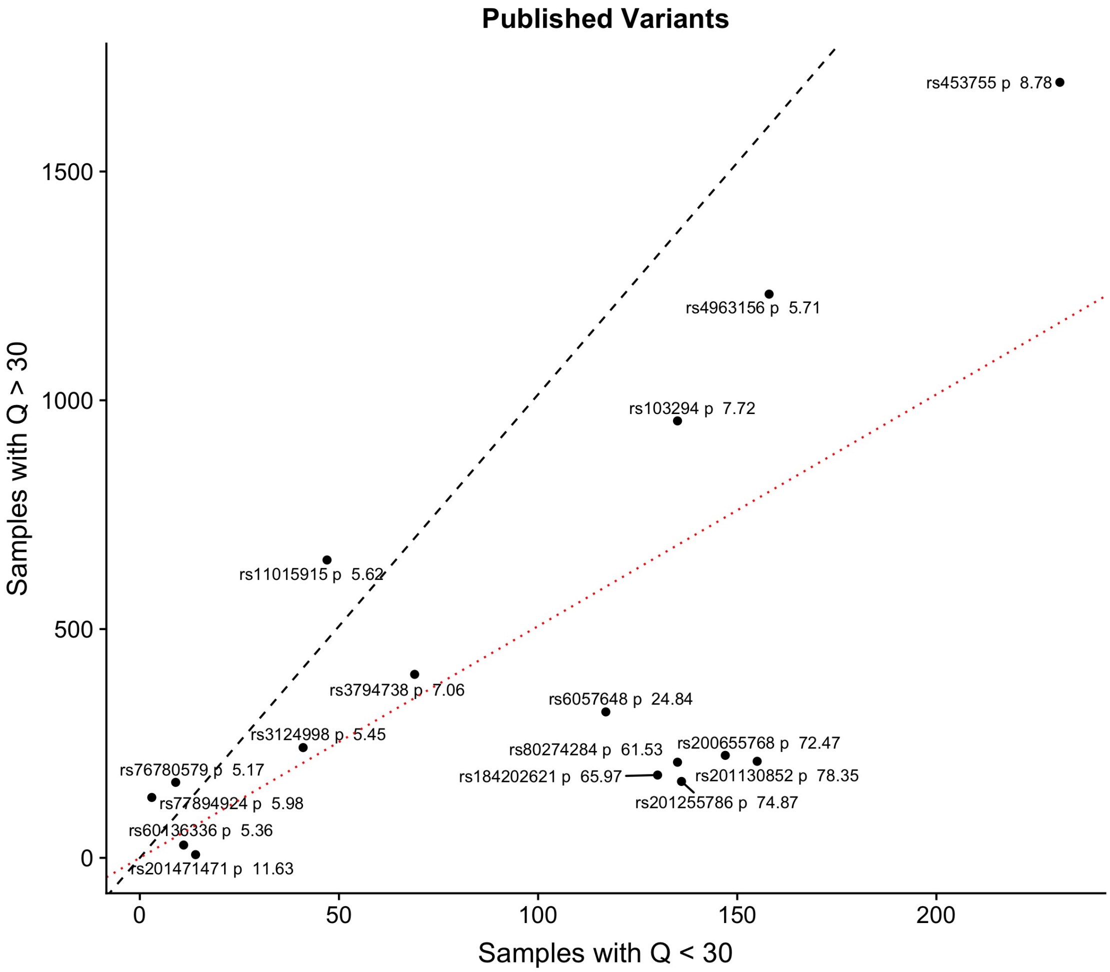


**Figure S21.** Site frequency spectrum plot comparing the frequency of $Q$-associated variants identified in publications, for individuals with $Q$ scores above and below 30. The black dashed lines indicates equal allele frequencies while the red dotted line for variants twice as frequent in individuals with $Q$ scores below 30. Each of the rsIDs of the variants are labelled for clarity.
